# Wild-type sTREM2 blocks Aβ aggregation and neurotoxicity, while the Alzheimer’s R47H mutant does the opposite

**DOI:** 10.1101/2020.12.03.409995

**Authors:** Anna Vilalta, Ye Zhou, Jean Sevalle, Jennifer K. Griffin, Kanayo Satoh, David H. Allendorf, Suman De, Mar Puigdellívol, Arturas Bruzas, Miguel A. Burguillos, Roger B. Dodd, Fusheng Chen, Yalun Zhang, Patrick Flagmeier, Lisa-Maria Needham, Masahiro Enomoto, Seema Qamar, James Henderson, Jochen Walter, Paul E. Fraser, David Klenerman, Steven F. Lee, Peter St George-Hyslop, Guy C. Brown

**Affiliations:** Department of Biochemistry, University of Cambridge, Tennis Court Road, Cambridge CB2 1QW, U.K; Departments of Medicine (Neurology) and Medical Biophysics, University of Toronto and University Health Network, Krembil Discovery Tower, 60 Leonard Avenue, Toronto, ON, M5T 0S8, Canada; Cambridge Institute for Medical Research, Keith Peters Building, Room 4.36, Cambridge Biomedical Campus, Hills Road, Cambridge CB2 0XY, U.K; Department of Chemistry, University of Cambridge, Lensfield Rd, Cambridge CB2 1EW, U.K; Princess Margaret Cancer Centre, University Health Network, Department of Medical Biophysics, University of Toronto, Toronto, ON, Canada; Molecular Cell Biology, Department of Neurology, University of Bonn, Venusberg Campus 1, 53105 Bonn, Germany

**Keywords:** Amyloid beta, Alzheimer’s disease, TREM2, microglia, oligomers, neurotoxicity, sTREM2

## Abstract

Missense mutations (e.g. R47H) of the microglial receptor TREM2 increase risk of Alzheimer’s disease (AD), and the soluble ectodomain of wild-type TREM2 (sTREM2) appears to protect in vivo, but the underlying mechanisms are unclear. We show that Aβ oligomers bind to TREM2, inducing shedding of sTREM2. Wild-type sTREM2 inhibits Aβ oligomerization, fibrillization and neurotoxicity, and disaggregates preformed Aβ oligomers and protofibrils. In contrast, the R47H AD-risk variant of sTREM2 is less able to bind and disaggregate oligomeric Aβ, but rather promotes Aβ protofibril formation and neurotoxicity. Thus, in addition to mediating phagocytosis, wild-type TREM2 may protect against amyloid pathology by Aβ-induced release of sTREM2 that blocks Aβ aggregation and neurotoxicity; while R47H sTREM2 promotes Aβ aggregation into neurotoxic forms, which may explain why the R47H variant gene increases AD risk several fold.

## Introduction

A prominent neuropathological feature of Alzheimer’s disease (AD) is the presence of extracellular deposits of the amyloid β-peptide in amyloid plaques, surrounded by activated microglia^1–3^. The importance of this microglial response to the pathogenesis of AD has been highlighted by the recent discovery of sequence variants in multiple genes expressed in microglia that alter risk for AD. Prominent amongst these microglial AD-risk genes is the “triggering receptor expressed on myeloid cells 2” (TREM2)^4–6^, of which there are several missense mutations in the ectodomain, including R47H, associated with increased risk for AD^4–6^. The biological mechanisms underlying this association remain unclear.

Full length TREM2 is expressed on the plasma membrane of microglia, where it can be cleaved by one or more metalloproteases to produce i) a membrane-bound C-terminal fragment (CTF); and ii) an N-terminal fragment consisting of the soluble ectodomain of TREM2 (sTREM2), which is released into the extracellular space^7–9^. sTREM2 has been thought of as a non-functional, degradation product of TREM2, and used as a biomarker of microglial activation^10–12^. However, several recent observations suggest the possibility that sTREM2 per se may play a role protecting against AD by interacting with Aβ. First, oligomeric Aβ binds to TREM2 and to sTREM2-Fc fusion protein^13–15^, suggesting that sTREM2 might bind Aβ and potentially affect its aggregation state. Second, injection or expression of sTREM2 in the hippocampus of 5xFAD mice reduces both amyloid plaque load and memory deficits^16^, indicating that sTREM2 inhibits amyloid pathology somehow. Third, in transgenic mouse models overproducing Aβ the knockout of TREM2 expression accelerates amyloid plaque seeding^17^ and the plaques have increased protofibrillar halos and hotspots^3,16,18^, indicating that either TREM2 or sTREM2 inhibit plaque formation. Fourth, in the earliest pre-symptomatic stages of AD, at the time of Aβ deposition, sTREM2 levels in cerebrospinal fluid (CSF) are lower than in healthy controls^10,19^, consistent with sTREM2 being an endogenous inhibitor of Aβ deposition. However, the CSF Aβ levels rise in the early symptomatic stages of AD, and then decline again at later stages of AD^11,12^. Fifth, mild cognitive impairment (MCI) and AD patients with higher sTREM2 levels in CSF have slower brain atrophy, cognitive decline and clinical decline^20,21^, consistent with sTREM2 inhibiting AD progression. Sixth, healthy controls and MCI patients with higher sTREM2 levels in CSF have slower progression of amyloid and tau deposition^22^, consistent with sTREM2 inhibiting Aβ aggregation and subsequent tau pathology.

All of these in vivo findings are compatible with the hypothesis that sTREM2 might protect against AD, potentially by impacting Aβ aggregation. To explore this hypothesis and the mechanisms involved, we investigated the interaction of Aβ with sTREM2 in vitro. We report that soluble Aβ oligomers bind TREM2 receptor on microglia and induce shedding of sTREM2. Next, we show that sTREM2 can bind and disaggregate Aβ oligomers, block Aβ fibrillization and reduce Aβ neurotoxicity. These activities are attenuated in the R47H TREM2 holoprotein and R47H sTREM2. Moreover, the R47H sTREM2 promotes formation of morphologically distinct Aβ protofibrils. These data indicate additional mechanisms by which TREM2 protects against Alzheimer’s disease, and a previously unrecognised mechanism by which the R47H mutant increases risk.

## Results

### Aβ oligomers bind TREM2 holoprotein and induce TREM2 ectodomain shedding

To confirm that TREM2 can act as a cell surface receptor for Aβ oligomers, we treated mouse primary microglia or TREM2-transfected HeLa cells with Aβ42 oligomers that have been characterized both by electron microscopy and by oligomer/fibril specific antibodies (Supplementary Fig. 1; note only Aβ42 was used in this work and will be referred to as Aβ henceforth). We then immunoprecipitated TREM2 from lysates of these cells, and found that Aβ co-immunoprecipitated with endogenous TREM2 on primary microglia from wild-type mice, but not on microglia from TREM2^−/−^ knockout mice (Supplementary Fig. 2i). The Aβ binding was prevented by TREM2-blocking antibody (Supplementary Fig. 2ii). Oligomeric Aβ bound to TREM2 but not TREM1 (Supplementary Fig. 2iii). Monomeric Aβ co-immunoprecipitated with TREM2 less efficiently than oligomeric Aβ (Supplementary Fig. 2iv and v, p = 0.03). The TREM2: Aβ oligomer interaction is at least partially specific because neither of two other neurodegeneration-associated oligomeric proteins (oligomeric α-synuclein or oligomeric Tau) bound to TREM2 even in HeLa cells overexpressing TREM2 and DAP12 (Supplementary Fig. 3). These results, confirm and extend the work of other groups showing that oligomeric Aβ binds TREM2^13–15^.

We next tested the consequences of Aβ oligomers binding to TREM2 on primary microglia. Aβ oligomers induced release of sTREM2 into the medium of primary microglia from wild-type mice, but not microglia from mice engineered to express R47H TREM2 (Supplementary Fig. 4i, p<0.05). To study this effect in more detail, we stably expressed TREM2 (together with DAP12) in HEK293 cells. Treatment of these cells with Aβ oligomers resulted in dose-dependent release of sTREM2 into the medium and the accumulation of TREM2-CTF in cell lysates (Figures 1i & ii & Supplementary Fig. 4ii). In contrast, treatment with Aβ monomers or fibrils resulted in no modulation of sTREM2’s release (Fig. 1iii & Supplementary Fig. 5). The endoproteolysis of TREM2 induced by oligomeric Aβ was attenuated in HEK293 cells expressing R47H TREM2 (Fig. 1ii & Supplementary Fig. 4ii & Fig. 5). Thus, oligomeric Aβ stimulates shedding of sTREM2 from wild-type TREM2, but less so from R47H TREM2.

**FIGURE 1.**
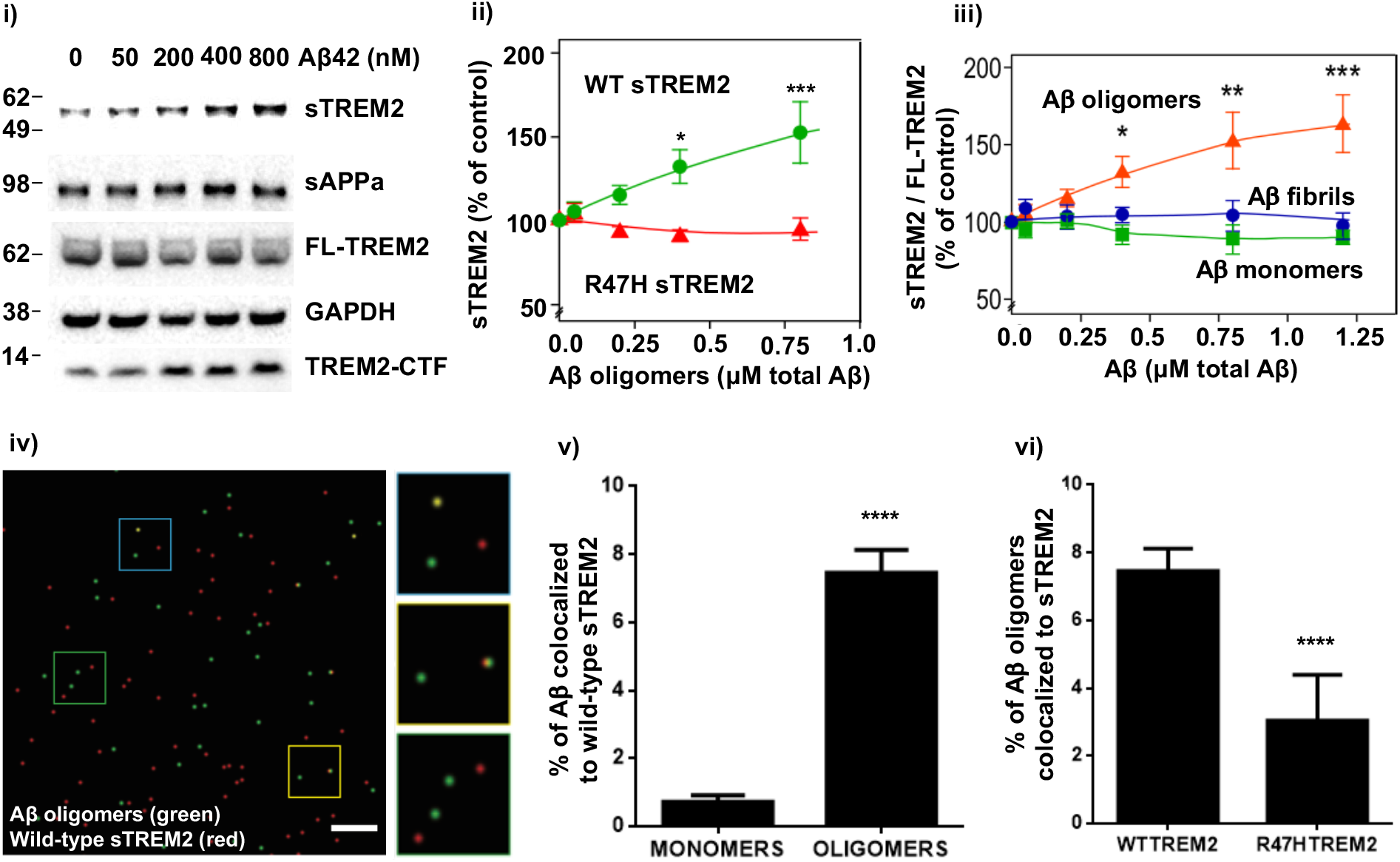
Aβ oligomers induce TREM2 proteolysis and sTREM2 release, which then binds Aβ oligomers, but R47H sTREM2 binds less. **i)** Western blot of cell lysate and supernatant (sTREM2) of HEK293 cells co-expressing DAP12 and full-length TREM2 (FL-TREM2)16 hours after adding Aβ oligomers. ii) Quantification of sTREM2 release from wild-type (green line) and R47H TREM2 (red line) expressing HEK293 cells. iii) Quantification of sTREM2 release from wild-type TREM2 expressing HEK293 cells induced by doses of Aβ oligomers (red line), monomers (green line) or fibrils (blue line). For both ii & iii) error bars = SEM; *= p<0.05 **=p<0.01 ***=p<0.001, n=3 independent experiments; one-way ANOVA with Tukey’s post-hoc multiple comparisons test. **iv)** Example field of single-molecule TIRF imaging of mixture of Aβ oligomers (green) and wild-type TREM2 ectodomain (red), where co-localised spots appear yellow. Scale bar: 1 micron. Magnified image of three sections of field at right. **v)** Proportion of monomeric or oligomeric Aβ colocalized with wild-type sTREM2. **vi)** Proportion of Aβ oligomers colocalized with wild-type or R47H TREM2 ectodomain. For v) & vi), error bars = SEM; ****p<0.0001, n=3 independent preparations, each analysed in 9 fields each; two tailed t test of significance.

### Wild-type sTREM2 binds Aβ oligomers better than R47H TREM2

Because Aβ oligomers bind to TREM2 and induce shedding of sTREM2, it is of interest whether sTREM2 itself binds Aβ oligomers. Three groups reported that wild-type sTREM2 fused to the Fc domain of IgG binds oligomeric Aβ^13–15^. However, Zhao *et al*^13^ and Zhong *et al*^14^ found that R47H sTREM2-Fc bound less than wild-type to Aβ, while Lessard *et al*^15^ found that they bound equally. We applied three orthogonal assays using an approach that did not require fusion of sTREM2 to the Fc domain of IgG, to reassess this question. Firstly, we examined this interaction at the molecular level using single molecule **<u>t</u>**otal **<u>i</u>**nternal **<u>r</u>**eflection **<u>f</u>**luorescence (TIRF) microscopy-based two-colour coincidence detection^23^. HiLyte 488-dye labelled Aβ was oligomerized and TAMRA-dye labelled sTREM2 added and imaged. We took advantage of the fact that monomeric Aβ bleaches rapidly, so co-localization of Aβ and sTREM2 before and after bleaching enabled us to distinguish between monomers and oligomers. These experiments revealed that wild-type sTREM2 bound oligomeric Aβ much more than monomeric Aβ (Fig. 1iv & v), and that R47H sTREM2 bound less well to oligomeric Aβ than wild-type sTREM2 did (Fig. 1vi).

Second, this result was confirmed by a semi-quantitative dot blot assay, which showed that wild-type sTREM2 preferentially bound oligomeric Aβ over monomeric Aβ, and that R47H sTREM2 bound less of both forms of Aβ (Supplementary Fig. 6i & ii).

Third, we used Bio-Layer Interferometry (BLI)^24^ to quantitatively assess the Aβ oligomer: sTREM2 interaction by immobilizing biotin-Aβ oligomers on Streptavidin biosensors and then exposing them to sTREM2. These studies showed that sTREM2 associated with Aβ oligomers with two different rates, and dissociated from Aβ oligomers with two different rates (Supplementary Fig. 6iii & iv). Best fit association and dissociation curves for both wild-type and R47H sTREM2 were with a 2:1 heterogeneous ligand model. In agreement with the semiquantitative experiments described above, these BLI studies confirmed that R47H sTREM2 bound Aβ oligomers less well than wild-type sTREM2 (Supplementary Table 1: **TREM2 WT** K_D_1 = 2.00 ± 0.15 μM, K_D_2 = 0.29 ± 0.08 μM; **TREM2 R47H** mutant K_D_1 = 11.70 ± 5.97 μM, K_D_2 = 1.22 ± 0.30 μM; where K_D_1 is the dissociation constant of the predominant interaction within the heterogeneous population of Aβ species).

### Wild-type sTREM2 inhibits Aβ oligomerisation and dissolves Aβ oligomers, whereas R47H sTREM2 induces Aβ protofibrils

Because sTREM2 bound to Aβ oligomers, we next investigated whether sTREM2 affected Aβ oligomerization. We incubated 22 μM monomeric Aβ ± 0.22 or 0.44 μM sTREM2 (wild-type or R47H) for 3 hours at 37°C, i.e. conditions known to generate Aβ oligomers. The formation of Aβ oligomers was then assessed by transmission electron microscopy (TEM) (Fig. 2, Supplemental Fig. 7; dot blotting with A11 anti-oligomer antibody; and by high performance liquid chromatography size exclusion chromatography (HPLC-SEC) experiments (Supplementary Figure 8). These experiments revealed that pre-incubation of Aβ monomers with wild-type sTREM2 (at molar ratios of 1:50 and 1:100 sTREM2: Aβ) inhibited Aβ oligomerisation (Fig. 2i) quantified by area (Fig. 2ii) or number (Supplementary Fig. 7i). Size exclusion chromatography and antibody dot blots confirmed that sTREM2 inhibited formation of Aβ oligomers (Supplementary Fig. 8i). Analysis of the size distribution of Aβ oligomers by TEM revealed that wild-type sTREM2 reduced the abundance of small oligomers (oligomer size <60 nm^2^), and eliminated larger oligomers (oligomer size >70 nm^2^) (Supplementary Fig. 9i).

In contrast, R47H sTREM2 did not block Aβ oligomerization, but instead promoted the formation of morphologically-distinct Aβ protofibrils (Figure 2iii & iv & Supplemental Fig. 7ii & 9ii). Thus, the presence of wild-type sTREM2 reduces the formation of Aβ oligomers, particularly larger oligomers; and in stark contrast, R47H sTREM2 induces Aβ monomers to form Aβ protofibrils.

**Figure 2.**
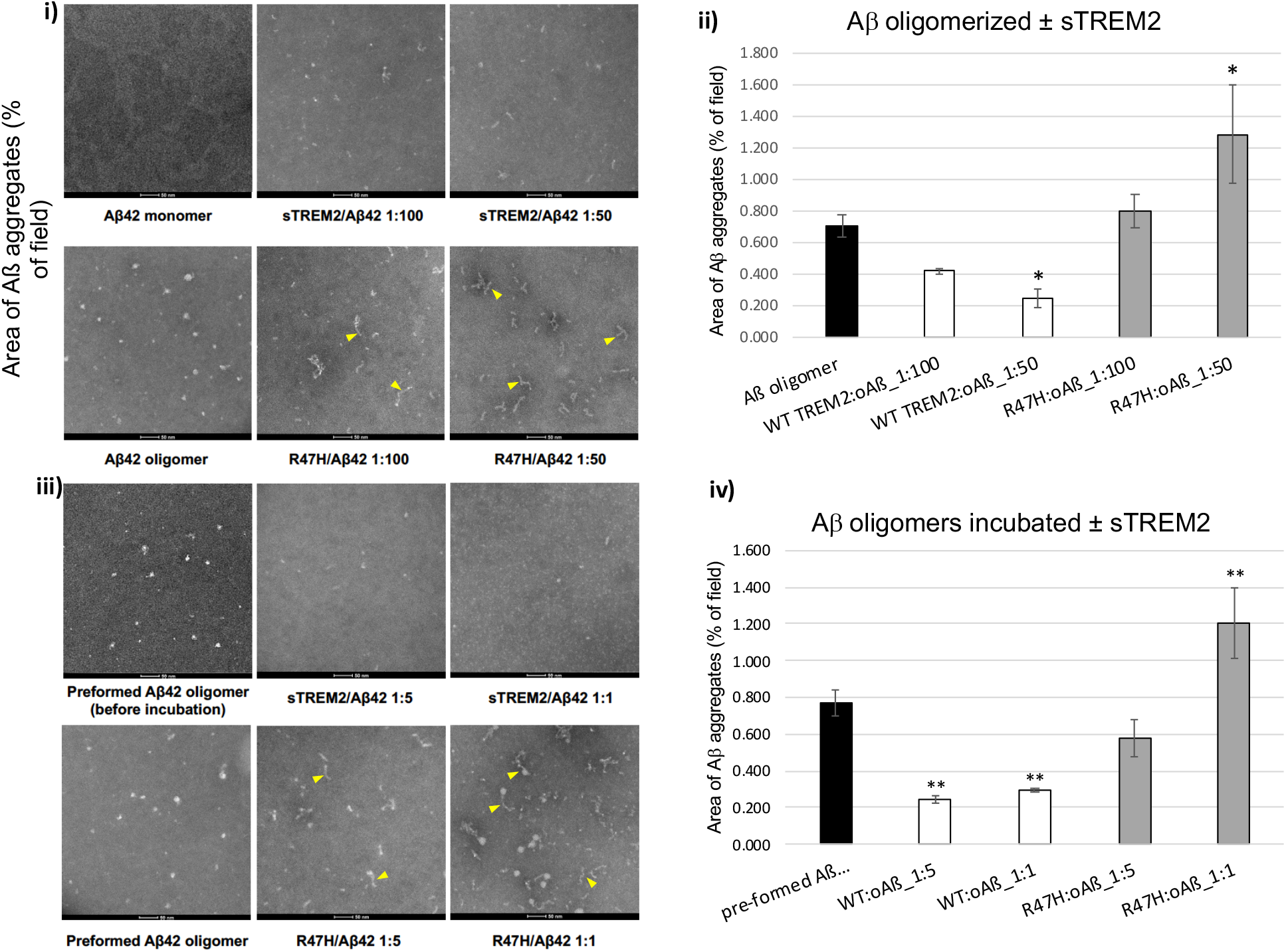
Wild-type sTREM2 inhibits Aβ oligomerization and disaggregates Aβ oligomers, whereas R47H sTREM2 converts Aβ to protofibrils. **i**) Aβ was aggregated for 3hr at 37°C ± WT sTREM2 or R47H at molar ratios of 1:100 or 1:50 (sTREM2: Aβ). TEM images revealed that WT sTREM2 reduced Aβ oligomerization, whereas R47H sTREM2 increased proto-fibrils (indicated by yellow arrowheads) and amorphous oligomers. **ii)** Quantification of three experiments indicated WT sTREM2 decreased and R47H increased area of Aβ aggregates. **iii)** Preformed Aβ oligomers were treated with WT sTREM2 or R47H at molar ratios of 1:5 or 1:1 for 30 min. TEM images revealed that preformed Aβ oligomers were disaggregated by WT sTREM2, but formed more globular Aβ oligomers and proto-fibrils with R47H sTREM2. iv) Quantification confirmed decreased area of Aβ aggregates with WT sTREM2, and increased area with R47H sTREM2. For both ii) & **iv)**, error bars represent SD, and statistical analysis was performed using one-way ANOVA followed by Bonferroni’s multiple comparison test (n=3 for each, \**p*<0.05 vs Aβ oligomer, \*\**p*<0.01 vs pre-formed Aβ oligomers).

To study reversal of oligomerization, pre-assembled Aβ oligomers were diluted to 2 μM and mixed with wild-type or R47H sTREM2 (at molar ratios of sTREM2: total Aβ of: 1:5 and 1:1) and incubated for 30 minutes at 37°C. The Aβ assemblies were then assessed using the same methods as above. This experiment revealed that wild-type sTREM2 disaggregated pre-formed Aβ oligomers (TEM in Fig. 2iii & iv & Supplementary Fig. 7ii; anti-A11 dot blot and HPLC-SEC assays in Supplemental Figures 8ii). Wild-type sTREM2 strongly reduced the abundance of all but the very smallest oligomers (Fig 2iii & Supplementary Fig. 9ii). In contrast, R47H sTREM2 increased the abundance of Aβ oligomers, and induced the formation of morphologically-distinct Aβ protofibrils (Figure 2iii & Supplementary Fig. 7ii & 9ii). Aβ protofibrils are large, linear Aβ oligomers, formed from Aβ monomers in a variety of conditions^25,26^, and are observed in CSF of MCI and AD patients, but not healthy controls^27^.

We repeated these experiments at lower concentrations, using preformed Aβ oligomers at 100 nM Aβ (monomer equivalent) incubated with 20 or 100 nM sTREM2. Wild-type sTREM2 (at either concentration) induced rapid dissolution of the Aβ oligomers (Supplementary Figs. 10 & 11). In contrast, treatment with R47H sTREM2 induced aggregation of Aβ oligomers into much larger Aβ assemblies (Supplementary Figs. 10 & 11).

### Wild-type sTREM2 inhibited and reversed Aβ fibrillization, whereas R47H was less effective

Because wild-type sTREM2 inhibited Aβ oligomerization, and disaggregated Aβ oligomers, we decided to test whether sTREM2 affected Aβ fibrillization. In the absence of sTREM2, Aβ (10 μM) fibrillized with standard lag-phase kinetics as measured with Thioflavin T fluorescence. However, 1 μM of wild-type sTREM2 almost completely prevented Aβ fibrillization (Fig. 3i–iii). 1 μM of R47H sTREM2 did not prevent Aβ fibrillization, but the Thioflavin T fluorescence was lower (Fig. 3i), suggesting fibrils with less β sheet structure. Repeating these experiments, at lower concentrations and higher temperature, gave similar results. Thus, fibrillization of 2 μM Aβ was substantially delayed by 20 nM sTREM2 (i.e. a 100:1 ratio), and prevented by 1 μM sTREM2 (Fig. 3iv). In contrast, at the same low dose (20nM), R47H sTREM2 had minimal effect on Aβ fibrillization (Fig. 3v).

**FIGURE 3.**
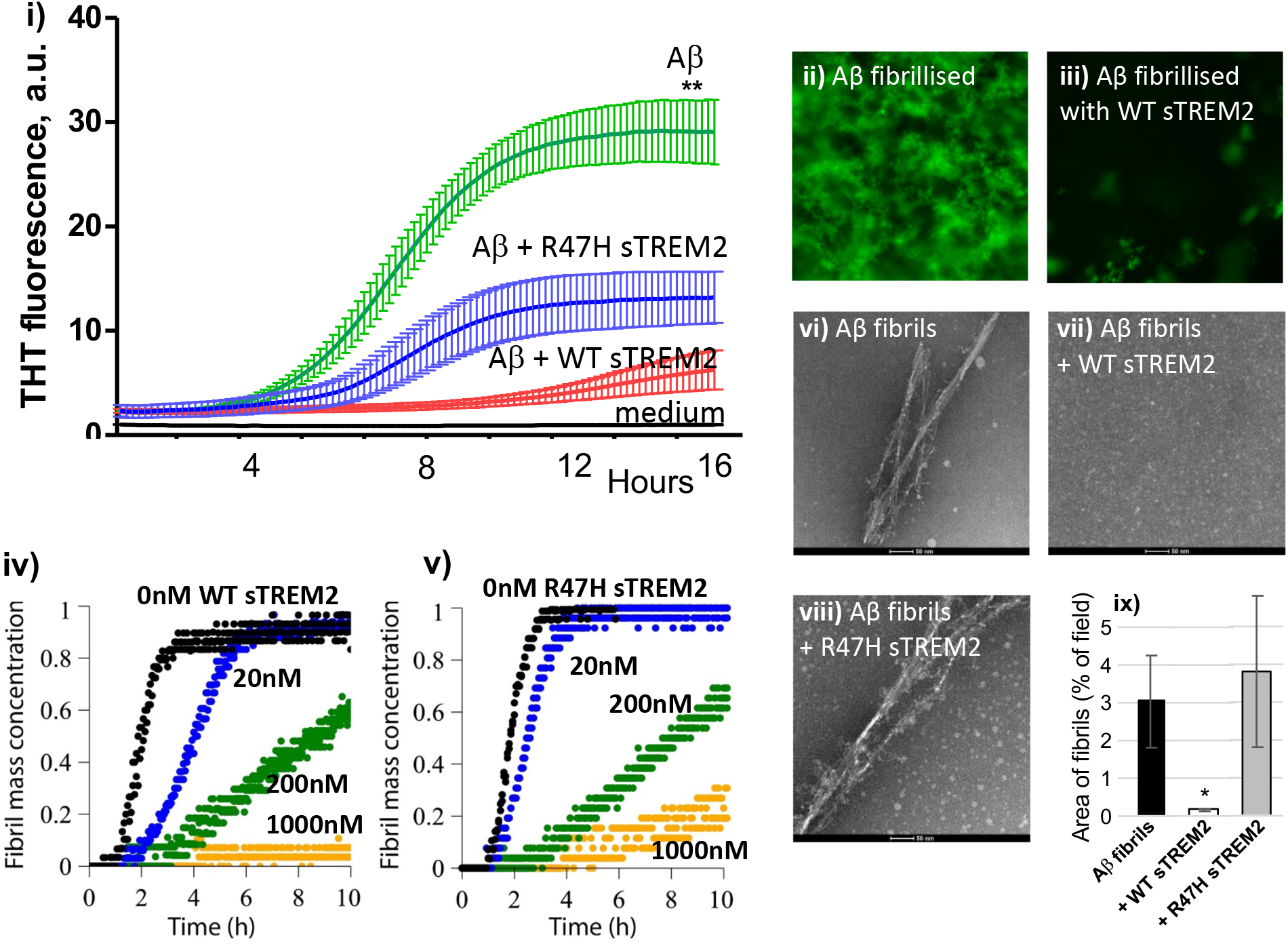
Wild-type sTREM2 blocks Aβ fibrillization, but R47H inhibits less. **i)** 10 μM of monomeric Aβ was incubated at 30°C in DMEM/F12 with 10 μM thioflavin T (THT) ± 1 μM wild-type or R47H sTREM2, and the fluorescence followed over time. Means and SD of 3 separate experiments are shown. There was significant difference (p<0.01, **) between the final fluorescence of the Aβ v Aβ + WT sTREM2 samples. At the end of the assay, **ii)** in the absence of sTREM2, and **iii)** presence of wild-type sTREM2, the bottom of the well was imaged using a fluorescence microscope with a x40 objective. **iv)** 2 μM of monomeric Aβ was incubated at 37°C in phosphate buffer with 10 μM THT + 0, 0.02, 0.2 or 1.0 μM wild-type sTREM2, and the fluorescence followed over time. **v)** 2 μM of monomeric Aβ was incubated at 37°C in phosphate buffer with 10 μM THT + 0, 0.02, 0.2 or 1.0 μM R47H sTREM2, and the fluorescence followed over time. Preformed Aβ fibrils were either **vi)** untreated, **vii)** treated with WT sTREM2, or **viii)** treated with R47H sTREM2 at molar ratios of 1:1 and TEM imaged after 30min. **ix)** Quantification of the area of Aβ fibrils confirmed that WT sTREM2 decrease Aβ fibrils, and R47H sTREM2 had no effect. Error bars represent SD. Statistical analysis was performed using one-way ANOVA followed by Bonferroni’s multiple comparison test (n=3-8, \**p*<0.05 vs pre-formed Aβ fibril).

To test whether sTREM2 could disaggregate fibrils of Aβ, small fibrils were pre-assembled and diluted to 2 μM (monomer equivalent) and mixed with wild-type or R47H sTREM2 (at molar ratios of sTREM2: total Aβ of 1:1) and incubated for 30 minutes, 2 hours or 24 hours at 37°C. Wild-type sTREM2 induced rapid and complete disaggregation of the preformed Aβ fibrils, whereas R47H sTREM2 had little effect on fibrils (Fig. 3vii–ix & Supplementary Fig. 12), although there may be an increase in oligomers (Fig. 3viii).

### Wild-type sTREM2 inhibits, while R47H sTREM2 increases Aβ oligomer neurotoxicity

Because sTREM2 bound Aβ oligomers and inhibited Aβ oligomerisation and fibrillization, we next tested whether sTREM2 affected the neurotoxicity of Aβ. Aβ neurotoxicity is thought to be mediated by 2 different mechanisms - namely by permeabilization of neuronal membranes^28^, and by microglial activation^28,29^.

Consequently, we initially used nanosized phospholipid membrane vesicles containing a calcium-sensitive fluorophore to explore the ability of Aβ to permeabilize membranes. Prior work has shown that Aβ oligomers (but not monomers or fibrils) can permeabilize these membranes 28. We found that wild-type sTREM2 and R47H sTREM2 proteins themselves had no effect on membrane permeabilization when added in the absence of Aβ (Supplementary Fig. 13i). By contrast, Aβ oligomerized for 6 hours in the absence of sTREM2 induced permeabilization of the vesicles. However, Aβ oligomerized in the presence of wild-type sTREM2 (at molar ratios of sTREM2: total Aβ of 1:10) induced significantly less permeabilization (Figure 4i). Wild-type sTREM2 also inhibited the permeabilization induced by pre-formed Aβ oligomers (at a 1:1 molar ratio) (Fig. 4ii). The simplest explanation for this inhibition of permeabilization is our previous finding that wild-type sTREM2 inhibits Aβ oligomerization and disaggregates Aβ oligomers at these concentrations. However, we do not discount the possibility that binding of wild-type sTREM2 to Aβ oligomers might also reduce their toxicity.

**FIGURE 4.**
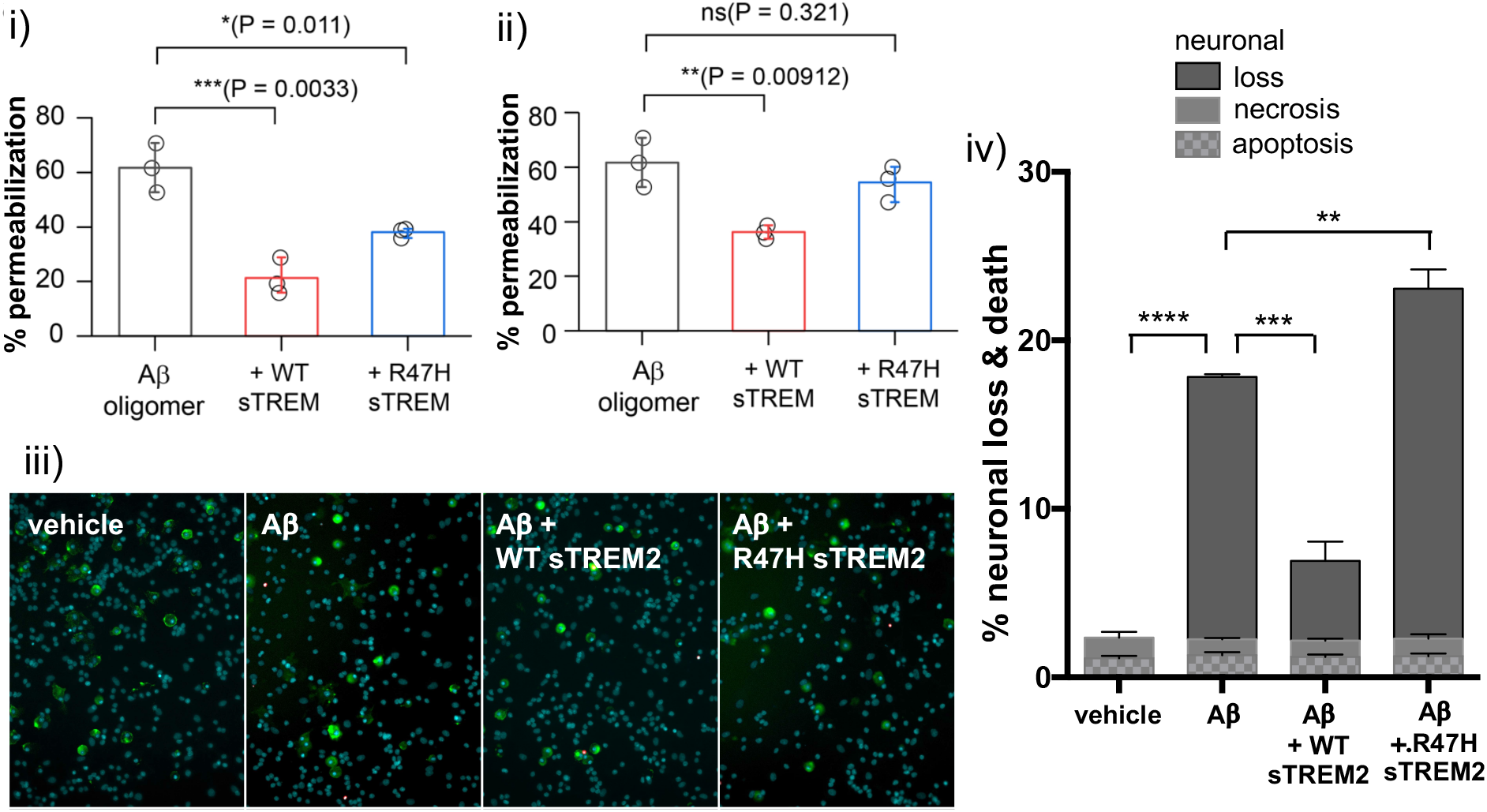
Wild-type sTREM2 blocks Aβ-induced membrane permeabilization and neuronal loss, but R47H sTREM2 inhibits less. **i)** 10 μM of monomeric Aβ was aggregated ± 1 μM wild-type (WT) or R47H sTREM2 for 6 hours, diluted to 200 nM and membrane permeabilization assay was performed. Error bars = SD of three independent experiments; **=p<0.01; statistics: two sample t-test. **ii)** 10 μM of monomeric Aβ was aggregated for 6 hours, diluted to 200 nM, then incubated for 30 minutes with vehicle, 200 nM WT sTREM2 or R47H sTREM2, before the membrane permeabilization assay was performed. Error bars = SD of three independent experiments; **=p<0.01; statistics: two sample t-test. **iii & iv)** Mixed neuronal-glial co-cultures were treated with either: vehicle, 250 nM monomeric Aβ, 250 nM Aβ + 25 nM WT sTREM2 or 250 nM Aβ + 25 nM R47H sTREM2. Three days later, cultures were stained with isolectin B4 (green, to identify microglia), propidium iodide (red, to identify dead cells) and Hoechst 33342 (blue, to identify cells and whether apoptotic) and imaged (representative fields: **iii)**, and the number of apoptotic, necrotic and healthy neurons were quantified (mean data: iv). Loss is the decrease in neuronal density relative to vehicle treated cultures. Error bars = SEM; **=p<0.01, ***=p<0.001, ****=p<0.0001; n=4 independent experiments on separate cell cultures. Statistical analysis was by one-way ANOVA and Tukey’s post hoc test.

When these experiments were repeated using R47H sTREM2 (at a molar ratio of sTREM2: total Aβ of 1:10) we found that Aβ oligomers formed in the presence of R47H sTREM2 also induced less permeabilization than Aβ oligomerized alone, but allowed more permeabilization than Aβ oligomerized with wild-type sTREM2 (Fig. 4i). In contrast to wild-type sTREM2, R47H sTREM2 did not significantly inhibit permeabilization induced by pre-formed Aβ oligomers (Figure 4ii). These results are consistent with wild-type sTREM2 reducing Aβ oligomer abundance, but R47H sTREM2 not reducing Aβ oligomer abundance (Fig. 2).

We next tested whether sTREM2 could block the neurotoxicity induced by Aβ in neuronal-glial co-cultures. We have previously shown that 250 nM Aβ induces slow, progressive neuronal loss mediated by microglia in this co-culture system^29^. We found that 250 nM Aβ induced neuronal loss over three days, and this Aβ-induced neuronal loss was substantially reduced by co-treatment with 25 nM wild-type sTREM2 (Fig. 4iii & iv). In contrast, co-treatment of the cultures with 25 nM R47H sTREM2 increased the neuronal loss above the level induced by 250 nM Aβ alone (Fig. 4iv). Wild-type and R47H sTREM2 proteins by themselves had no significant effect on neuronal loss (Supplementary Fig. 13ii). As membrane permeabilization is thought to be mediated by smaller Aβ oligomers, while microglial activation is thought to be mediated by larger Aβ aggregates^28^, our finding that R47H sTREM2 increased Aβ-induced neuronal loss (Fig. 3iv), which is known to be microglia-mediated^29^, is consistent with R47H sTREM2 increasing larger Aβ aggregates (Fig. 2). Overall, our results indicate that wild-type sTREM2 reduces Aβ neurotoxicity, while R47H TREM2 increases Aβ neurotoxicity.

## Discussion

We have shown that Aβ oligomers, but not Aβ monomers or fibrils, bind to microglial TREM2 and induce sTREM2 release, and this released sTREM2 also preferentially binds to Aβ oligomers, consistent with previous reports^13–15^. Crucially however, we also found that wild-type sTREM2 inhibits the formation of fibrils and larger Aβ oligomers and disaggregates protofibrils and larger Aβ oligomers into the smallest Aβ oligomers. In contrast, R47H sTREM2 promoted the formation of Aβ protofibrils, indicating a gain of function by this mutation. These effects may provide a partial explanation of how a single copy of the TREM2 R47H mutant is associated with increased risk for AD, by increasing production of neurotoxic forms of Aβ.

One potential caveat is the protein concentrations used in our experiments. In order to observe Aβ aggregation over a reasonable experimental timeframe, the lowest concentration of Aβ used here was 100 nM, whereas levels found in CSF are about 0.1 nM ^10,20,30^. However, the Aβ oligomer concentrations near amyloid plaques was estimated as 700 nM^31^, and plagues are likely be a more relevant location for Aβ aggregation in AD than CSF. sTREM2 concentrations in lumbar CSF average about 0.2 nM^10,20,30^; however again, it is likely that sTREM2 levels are higher near amyloid plaques surrounded by activated microglia, which express high levels of TREM2^3^. Note also that the observed effects of sTREM2 on Aβ are rapid: 20 nM sTREM2 dissolved 100 nM Aβ oligomers within 30 mins in vitro (Supplementary Fig 11). Lower concentrations are likely to have the same disaggregating effect over a slower time course, which may be more relevant to the slow time course of AD.

We did not investigate the molecular mechanisms by which sTREM2 affected Aβ aggregation, but our finding that wildtype sTREM2 stabilises the smallest observable oligomers (Supplementary Figure 9 & 11), suggests the possibility that sTREM2 binds preferentially to these (or undetectable oligomers such as dimers), thereby potentially blocking further growth of aggregates, and disaggregating larger Aβ aggregates into smaller, potentially non-toxic forms. Precedent for this is the finding that α-synuclein-specific single-domain antibodies (nanobodies) bind α-synuclein stable oligomers and convert them into less stable oligomers with reduced toxicity^32^. However, clearly more research is required to determine the mechanisms for sTREM2 and Aβ.

In summary, our experimental data reveal how Aβ oligomer binding to TREM2 may mediate direct (via activation of intracellular TREM2 dependent signalling pathways) and indirect protective mechanisms (via effects on Aβ oligomer assembly and toxicity). Our studies are broadly congruent with previous research showing that knockout of TREM2 expression in APP mice, resulted in accelerated amyloid plaque seeding^17^, with increased protofibrillar halos and hotspots around these plaques^3,16,18^. Our studies provide a potential explanation of the clinical observation of slower rates of cognitive and clinical decline in patients with MCI or AD who have higher levels of sTREM2 in the CSF^20,21^, and slower rates of amyloid deposition in healthy controls and MCI patients with higher sTREM2 levels in CSF^22^. They also provide a potential explanation for the beneficial effects of infusing or expressing sTREM2 into the brain of mouse models of AD^16^. Additional biophysical studies will be required to identify the key Aβ species dynamically interacting with wild-type sTREM2 to prevent neurotoxicity, and the Aβ species stabilised by sTREM2 R47H to increase neurotoxicity. This knowledge could be exploited for the design of small brain-penetrant molecular mimics of sTREM2 as potential therapeutics for AD.

## METHODS

### cDNA constructs

Wild-type human TREM2 (hTREM2) and DAP12 (hDAP12) were subcloned in pCMV6-A vector (Origene, Rockville, MD, USA). A single C→A nucleotide polymorphism (SNP) at codon 85 (T85K) on the Origene TREM2 cDNA clone was reverted to consensus wild-type sequence by site-directed mutagenesis.

### Antibodies

The following antibodies were used: human TREM2 (#AF1828, 1:100 for immunocytochemistry (ICC); 1:1000 for western blot; R&D); mouse TREM2 (#AF1729, 1:500 for western blot; R&D); DDK (FLAG) (#TA50011, 1:2000 for western blot; Origene); V5 (#46-0705, 1:2000 for western blot; Life Technologies, Burlington); Nicastrin (#sc-14369, 1:200 for western blot; Santa Cruz); GAPDH (#2118, 1:3000 for western blot; Cell Signaling); Anti-His (C-term)-HRP (#46-0707, 1:2000 for dot blot, Invitrogen); Aβ (6E10, 1:1000 for western blot and dot blot, Covance); TrueBlot secondary antibodies conjugated with horseradish peroxidase (anti-goat IgG, anti-mouse IgG and anti-sheep IgG, 1:1000 for western blot after immunoprecipitation; Rockland); Secondary antibodies conjugated with horseradish peroxidase (anti-mouse IgG; anti-rabbit IgG, 1:2000 for western blot; Thermo Scientific). Function-blocking TREM2 antibody (Clone 78.18, 25 μg/ml for blocking assay, AbD Serotec catalogue number MCA4772EL) and a control antibody (Rat IgG1, AbD Serotec) were used for blocking experiments.

### Animals

All experiments were performed in accordance with Canadian Council on Animal Care guidelines and UK Animals (Scientific Procedures) Act (1986) and approved by the Animal Care Committee at the University of Toronto and the Cambridge University local ethics committee.

TREM2 deficient mice were obtained from Dr. J. Gommerman, and were constructed by targeted homologous recombination^33^, which removed Exon 1 and 2, which include the start codon and the major extracellular IgG domain. In contrast to the recently reported Velocigene construct, the direction of the Hygromycin cassette was “reversed”. Crucially, in agreement with two other models, but in contrast to the Velocigene construct, RT-PCR analyses confirmed specific loss of TREM2 expression without the perturbation of TREM1L expression observed in the Velocigene construct^33^.

### Primary microglial cell culture

Primary cultures of mixed glial cells and pure microglia were prepared from the cerebral cortex of P0-3 day old C57BL/6 TREM2^+/+^ or TREM2^−/−^ mice as described^33^. After removal of olfactory bulbs, cerebral hemispheres were cut into ~1mm pieces, vortexed for 1 minute, filtered through a 40 μm cell strainer and plated in 80 cm^2^ tissue culture flasks. Mixed glia were cultured until microglia appeared (10-18 days after plating). Microglia were isolated by shaking flasks. Purified microglia were replated for analysis. On average, each experiment used microglia from 30-35 pups. All pups used in this study (wild type; R47H, and TREM2KO) were maintained on a C57BL/6 background.

### Mixed neuronal-glial co-cultures and treatments

Mixed neuronal-glial co-cultures were prepared from cerebella of 5-7 day-old rats^33^, cultured for 7 days, then treated with either: vehicle, 250 nM monomeric amyloid beta 1-42 (Aβ, 250 nM Aβ + 25 nM wild-type sTREM2 or 250 nM Aβ + 25 nM R47H sTREM2 for 3 days. Three days later, neuronal death and loss was quantified by staining the live cultures with propidium iodide for necrosis, with isolectin IB4 for microglia, and with Hoechst 33342 for nuclei, which is used to distinguish healthy (uncondensed) from apoptotic (condensed) nuclei, and to distinguish astrocytes (large bean-shaped nuclei) from neurons and microglia (small, round nuclei). The number of apoptotic, necrotic and healthy neurons, astrocytes and microglia are then counted on photos of multiple set fields, as previously described^29,33^,

### Amyloid β-peptide

Human Aβ42 (used in TREM2 cleavage), HiLyte Fluor 488-labelled Aβ42 (used in Single molecule TIRF imaging), HiLyte Fluor 647-labelled Aβ42 (used in IP and dot blot) and Biotin-Aβ42 (used in BLI) were purchased from AnaSpec and Bachem; Aβ42 oligomers were prepared as endotoxin-free preparations^34^, and validated by antibodies, gels and electron microscopy (Supplimentary Figure 1A). Prior work has established that these fluorescent tags do not significantly impact Aβ assembly^35^.

### Recombinant ectodomain expression and purification

cDNAs encoding residues 19-143 of WT and R47H TREM2 were ordered as linear DNA strings (Life Technologies) and inserted into pHLSec. The plasmids were transfected into Freestyle 293-F cells (Life Technologies) grown in suspension in FreeStyle293 medium. Cells were transfected at a density of 2.5 × 10^6^ / ml with a DNA concentration of 3 μg/ml and polyethyleneimine (Linear PEI 25 kDa molecular weight, PolySciences Inc) at 9 μg/ml. On day 6 post-transfection, conditioned medium was harvested and TREM2 protein was purified using IMAC (Histrap excel, GE Healthcare) followed by gel filtration chromatography in storage buffer - 20 mM HEPES, 200 mM NaCl, pH 7.0. To maintain a low endotoxin level: endotoxin-free chemicals and plasticware were used; all chromatography media, columns and concentrators (Vivaspin Turbo) were soaked for > 12 hours in 1 M NaOH. Proteins in storage buffer were concentrated to between 5-10 mg/ml for use and tested for endotoxin with the EndoZyme recombinant Factor C assay (Hyglos GmbH). Endotoxin levels in all assays were maintained at < 0.1 EU/ml.

### Co-immunoprecipitation and western blot

Primary microglia were rinsed with ice-cold DMEM without phenol red and incubated with 100 nM HiLyte Fluor 647-labelled Aβ at 4°C for 1 hour. For antibody blocking assays, before Aβ incubation, microglia cultures were pre-incubated with 25 μg/ml monoclonal anti-TREM2 antibody or rat IgG1 control for 20 minutes Cells were collected by scraping, washed with ice cold 1xPBS and centrifuged. Primary microglia were lysed with 1% CHAPSO buffer (1% 3-[(3-cholamydopropyl) dimethylammonio]-2-hydroxy-1-propanesulfonate (CHAPSO), 25 mM HEPES, 150 mM NaCl, 2 mM EDTA, pH 7.4). Crude membrane fractions were extracted from HeLa cell pellets as described^36^ and lysed with 1% CHAPSO buffer. Primary antibodies and corresponding IgG controls (1 μg/reaction) were incubated with lysates at 4°C for 2 hours, then mixed with Protein G Dynabeads (Life Technologies), incubated at 23°C for 20 minutes, then washed and eluted as per manufacturer’s instructions. The immunoprecipitates were separated on 4-12% MES/Bis-Tris gels (Life Technologies). Fluorescent signals of Aβ were detected using an Odyssey FC imaging system (LI-COR). Proteins were transferred onto nitrocellulose membranes, which were probed with primary antibodies and corresponding TrueBlot secondary antibodies.

In vitro co-precipitation of TREM2 or TREM1 with Aβ oligomers was performed by mixing recombinant Fc-tagged TREM2 or TREM1 ectodomain (residues 19-171; 100 ng/ml, R&D systems), 100 nM HiLyte Fluor 647-labelled Aβ oligomers and Protein G Dynabeads in 0.5% fatty acid free BSA PBS-Tween buffer. The mixture was incubated at room temperature for 1 hour and then washed, eluted and analysed the same as described above.

### Dot blot

Aβ monomers or oligomers (1 μg/dot) prepared as described above and anti-TREM2 antibody (0.3 μg/dot) were spotted onto a nitrocellulose membrane. Membrane strips were blocked with 3% fatty acid free BSA (Sigma Aldrich) and then incubated with 100 nM recombinant His-tagged WT/R47H TREM2 ECD diluted in blocking buffer at 4°C for 1 hour. Bound TREM2 ECDs were probed with HRP-conjugated anti-His tag antibody and detected with Odyssey FC imaging system.

### Bio-Layer Interferometory

#### i) Preparation of Aβ oligomers for Bio-Layer Interferometry (BLI) studies

Synthetic human Aβ residues 1-42 and biotinylated Aβ42 were purchased from Bachem or AnaSpec and solubilized in hexafluoroisopropanol (HFIP) with DMSO (200 μl) to generate monomeric species. Solubilized Aβ was diluted in filtered assay buffer [20 mM HEPES, 500 mM NaCl pH7.4] containing DMSO (2%) to a total volume of 9800 μl and a peptide concentration of 0.1 mg/ml (22 μM). Peptide solutions were aliquoted and incubated for 3 hours at 37°C with stock solutions stored at −80°C. Transmission electron microscopy (TEM) was performed using undiluted biotinylated Aβ (100%) or a mixture of biotinylated Aβ (10%) and unlabelled Aβ (90%) which were diluted to final concentrations of 1.1μM (Supplementary Figure 1iii & iv). These TEM samples (10 μl) were applied to carbonate coated grids and negatively stained with 1% phosphotungstic acid (PTA). TEM micrographs indicating the globular morphology of the Aβ oligomers were obtained on a Hitachi H-7000 operated at 75kV. The biotinylated oligomers were indistinguishable from non-biotinylated Aβ oligomers.

#### ii) Binding affinity of TREM2 ectodomain to Aβ was analyzed by Bio-Layer Interferometry (BLI) using the Octet RED384 system (ForteBio)

Assays were performed at 30°C in the assay buffer (20 mM HEPES, 500 mM NaCl, 0.1% w/v BSA, 0.02% v/v Tween20, pH 7.4) with vibration at 1,000 rpm. Streptavidin biosensors loaded with Biotin-Aβ (Anaspec) were incubated with TREM2 ectodomain WT (1, 2, 5, 10 μM) or R47H (2, 5, 10, 20 μM) to obtain the association curves, and subsequently in the assay buffer to obtain the dissociation curves. Kinetics data were analyzed using a 2:1 heterogeneous ligand model.

### Single molecule TIRF imaging

#### i) Aggregation of Aβ for TIRF studies

Stock solutions of HiLyte™ Fluor 488- labeled Human Aβ42 (Aβ, 0.1 mg; AnaSpec, USA) were prepared by dissolving the lyophilized peptide (1% NH4OH, 50 μl) followed by dilution into pH 7.4 PBS to 200 μM, aliquoted and stored at −80°C. Working solutions were prepared by diluting the monomeric stock solutions into pH 7.4 PBS on ice to the concentration used for the aggregation (0.5 μM). The working solution was then placed into a shaking incubator (3°C, 200 rpm) for 6 hours to ensure a significant population of oligomeric species. All protein samples were stored and diluted into LoBind microcentrifuge tubes (Eppendorf, Hamburg, Germany).

#### ii) Preparation of slides for TIRF microscopy

Slides were prepared in the same way as previously described^23^. Slides containing unlabelled Aβ and TREM2 were tested for fluorescence artefacts.

#### iii) TIRF Microscope set-up

Co-localization experiments were performed on a bespoke TIRF set-up. Continuous wave solid-state lasers operating at 488 nm and 641 nm were used for imaging. The beam power was controlled by attenuation through a neutral density filter, following which undesirable wavelengths were filtered out through excitation filters (LL01-488-25, FF01-640/14-25, Semrock). The beams were then circularly polarised by a quarter wave plate. Laser lines were combined by dichroic mirrors. The beams were then passed through a Köhler lens into the back aperture of an inverted microscope body (Olympus, IX73) and were reflected by a dichroic mirror (Di01-405/488/532/635, Semrock) through an oil immersion objective (Olympus, 60XOTIRF). Emitted radiation was collected through the same objective and passed through the dichroic before being filtered (FF01-480/40-25, LP02-647RU-25, Semrock). The fluorescence emission was then projected onto an EMCCD camera (Evolve Delta, Photometrics), each pixel was 275 nm in length. Data visualisation achieved through Micro-Manager and ImageJ software.

#### iv) Single-molecule TIRF Imaging

The co-localization experiment was designed to investigate binding of Aβ oligomers with TREM2. The Aβ 6 hour aggregation contained a mixture of monomeric and oligomeric species hence illumination was performed for significant time periods to photo-bleach the monomeric species. Automated TIRF co-localization experiments were performed with the use of a custom script (BeanShell, micromanager). Each data set consisted of a 3 × 3 grid of 9 images at different areas of the coverslide, distances between images was 350 μm as previously described^20^. Images were recorded at 33 ms exposure, beginning with 800 frames excitation with 641 nm followed by 800 frames excitation 488 nm excitation in the same field of view.

#### v) Co-localization Analysis

Co-localization data was analyzed with a bespoke ImageJ macro. The 488 nm illumination channel contained a mixture of monomeric and oligomeric Aβ species, it was determined that the majority of monomeric species were photo-bleached by 40 frames of 488 nm illumination. Therefore, only frames after this point were considered in this analysis. The 488 nm and 641 nm illumination channels were compressed in time to create two single frame images representing the average pixel intensities. Following which, points of intensity representing HiLyte™ Fluor 488 Aβ or AF647-TREM2 above a background threshold were located, counted and binary images of these maxima were created. The two images were then summed to identify co-localized points. Chance co-incident spots were excluded by 90° rotation of the binary image representing AF647-TREM2 points and summation with the HiLyte™ Fluor 488 Aβ image. Chance coincident spots were subtracted from the actual coincidence value and percentage coincidence was calculated with the equation below:

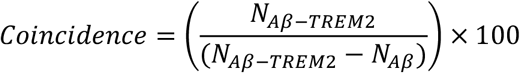

### TREM2 cleavage assays

Primary microglia from wild and TREM2 mutant mice were isolated from mixed glia culture by shaking and grown on 96-wells plates for 24 hours. Media were replaced by treatment media (1 ml of OptiMEM supplemented with 0.5% FBS) containing defined concentrations of freshly made Aβ oligomers. Cells were collected 24 hours after treatment and NTF release in conditioned media and TREM2 in cell lysate were detected by ELISA. TREM2 ELISA was performed on 96-well plates previously coated by TREM2 antibody (clone 78.18) and blocked with 0.5% BSA in 0.05% Tween20 in 1x PBS (PBST). Medium or cell lysate samples were added to pre-coated wells and incubated overnight at 4°C. After washing with PBST, wells were incubated with TREM2 antibody (sheep anti-mouse TREM2, AF1729) diluted in blocking buffer for 2 hours at room temperature. After secondary antibody (anti-sheep-HRP, 1:5000) incubation for 1 hour at room temperature and washing, TMB chromogenic substrate was added to each well and developed at room temperature in dark for 20 minutes. The absorbance at 450nm was read immediately after stop solution was added. TREM2 cleavage was represented by the ratio of relative values of sTREM2 in medium and total TREM2 in cell lysate, after subtracting the corresponding background values from TREM2 KO microglia in the same ELISA experiment.

### sTREM2 effects on Aβ oligomers

#### i) Preparation of Aβ oligomers and Aβ fibrils and sTREM2 treatment

HFIP-treated human synthetic Aβ42 were purchased from Bachem. Aβ solubilized by DMSO was diluted in PBS (pH 7.4) to a peptide concentration of 0.1 mg/mL (22 μM). To study sTREM2 effects on assembly of Aβ oligomers, monomeric Aβ (22 μM) solutions were incubated with or without WT sTREM2 or sTREM2 R47H (sTREM2: Aβ, 1:100 or 1:50) for 3 hours at 37°C. To study reversal of oligomerisation, monomeric Aβ (22 μM) solutions were incubated for 3 hours at 37°C to make oligomers then diluted and mixed with WT sTREM or sTREM2 R47H (sTREM2: Aß, 1:5 or 1:1) to a Aβ concentration of 2.2 μM. Then Aβ and sTREM2 mixture were incubated for 30 minutes at 37°C. To study dissociation of fibrils, monomeric Aβ (110 μM) solutions were incubated for 24 hours at 37°C with shaking (150 rpm) then diluted and mixed with WT or R47H sTREM2 (sTREM2: Aβ, 1:1) to a Aβ concentration of 2.2 μM, then incubated for 30 minutes, 2 hours or 24 hours at 37°C. These peptide solutions were then used for TEM, dot-blot and HPLC-SEC experiments.

#### ii) Transmission electron microscopy (TEM)

Aβ oligomers were prepared as described above. Transmission electron microscopy (TEM) was performed to characterize Aβ or mixtures of sTREM2 and Aβ which were diluted to final Aß concentrations of 1.1 μM or 100 nM. These TEM samples (10 μL) were applied to carbonate coated grids for 1 min and negatively stained with 1% phosphotungstic acid (PTA) for 1min. TEM micrographs were obtained on a Hitachi H-7000 operated at 120 kV.

#### iii) Dot blots

Aβ oligomers were prepared as described above. Aβ and mixtures of sTREM2 and Aβ oligomers (1 μg/dot) prepared as described above were spotted onto a nitrocellulose membrane. Membrane strips were blocked with 5% fatty acid free BSA (Sigma Aldrich) and then incubated with anti-oligomer specific antibody (A11, ThermoFisher Scientific) or anti-amyloid β sequence specific antibody (4G8, Millipore) for 1hr at RT and probed with HRP-conjugated anti-rabbit or mouse IgG antibodies and detected with ChemiDoc XRS+ imaging system (Bio-Rad).

#### iv) HPLC-SEC

Aβ oligomers were prepared as described above. The size of prepared Aβ peptides were analyzed by size exclusion chromatography (SEC). Samples were run on a BioSep SEC-S 4000 (Phenomenex) column using a ProStar 210 HPLC (Varian) system with a ProStar 325 UV detector (Varian). Injected samples were eluted with PBS (pH 7.4) at a flow rate of 0.5 mL/min at ambient temperature, and data were obtained at 280 nm. Peak areas were integrated using Star 6.2 Chromatography Workstation (Varian).

### Aβ fibrillization assay

10 μM of monomeric Aβ was incubated at 30°C in DMEM/F12 with 10 μM Thioflavin T ± 1 μM wild-type or R47H sTREM2 in Corning^®^ 96 Well White Polystyrene Microplate with a clear flat bottom in a FluoroSTAR Optima plate reader (with orbital shaking between readings) to monitor fluorescence. Alternatively, 2 μM of monomeric Aβ was incubated at 37 °C in 20 mM phosphate buffer (pH 8) with 10 μM Thioflavin T ± 0, 0.02, 0.2 or 1.0 μM wild-type or R47H sTREM2 in a fluorescence plate reader with a 440-nm excitation filter and a 480-nm emission filter without stirring as previously described^37^.

### Membrane permeabilization assay

Lipid vesicles were prepared as previously described^38^, using a 100:1 mixture of phospholipids 16:0-18:1 phosphatidylcholine and biotinylated lipids 18:1-12:0 Biotin phosphatidylcholine. The mean diameter of vesicles was ~ 200 nm and were filled with 100 μM Cal-520 dye using five freeze-and-thaw cycles. Non-incorporated Cal-520 dye was separated from the dye filled vesicles by size-exclusion chromatography. The vesicles were immobilized in biotin labelled PLL-g-PEG coated glass coverslips using a biotin-neutravidin linkage. Then coverslips were incubated with of 50 μL Ca^2+^ containing buffer solution and different position of the coverslips imaged (Fblank). Stage movements were controlled using an automated bean BeanShell-based program which allows fields-of-view selection without user bias. Then the 10 μM aggregation (6-hour time point) solution was added to the coverslips so that the final concentration of the peptide under the coverslip was 200 nM. Then the sample was incubated with the vesicles for 20 minutes and the same fields of view of each coverslip were imaged (Fsample). Then, 10 μL of 1 mg/mL of ionomycin was added to the same coverslips and the same fields of view were imaged again (Fionomycin). Ionomycin is a calcium ionophore and causes saturation of the fluorescence intensity observed. This allows us to normalise fluorescence originating from individual vesicles and subsequently quantify the sample induced Ca^2+^ influx using the following equation:

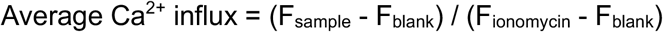

All the imaging was done using a Total Internal Reflection Fluorescence Microscope (TIRFM) based on an inverted Nikon Ti microscope. For excitation, a 488 nm laser beam was focused in the back focal plane of the 60x, 1.49 NA oil-immersion objective lens. Fluorescence signal emerged from the samples collected from the same objective and imaged air cooled EMCCD camera. All the imaging experiments were performed at 293K. Fluorescence images were acquired using a 488nm laser (~10 W/cm^2^) for 50 frames with a scan speed of 20 Hz and analyzed using ImageJ to calculate the localized fluorescence intensity of each vesicle.

### Statistics

All statistical analyses were performed using Prism (GraphPad Software Inc.). Non-parametric statistical tests were used when data did not show a normal distribution. The specific statistical test used in each experiment are described in the associated figure legend. Data are expressed as means ± SEM. Results were considered significant if p<0.05. In most cases, the sample size (i.e. the number of biological repeats, n) was 3, as indicated in the figure legends, and this number was selected as the default for experiments using isolated proteins, based on previous experience of variance sizes. In the case of Fig 3 ix, measuring the area of fibrils, the variance swas found to be high (because the number of fibrils is low and size large), so n was increased to 8 for Aβ alone, but remained at n=3 for Aβ + WT or R47H sTREM2. For Fig 4 iv, which is a primary cell culture experiment, the sample size was four, based on previous experience of variance sizes. The sample size n refers to biological repeats, which means repeating using a different biological sample preparation. Outliers were only encountered in the data of Fig 2 ii, and were identified and removed by using GraphPad Prism outlier calculator.

### Resource Availability

The published article includes all data generated or analyzed during this study. Original source data for Figures in the paper are available upon request to the corresponding authors. No proprietary software was used in the data analysis. Further information and requests for resources and reagents should be directed to

## Acknowledgements

GCB, JW & PStGH have received funding from the Innovative Medicines Initiative 2 Joint Undertaking under grant agreement No 115976 (PHAGO consortium), and from the Canadian Institute of Health Research (PStGH & PEF), Canadian Consortium on Neurodegeneration in Aging, Wellcome Trust, Medical Research Council, Zenith Award Alzheimer Association (PStGH).

## Author Contributions

AV, YZ, JS, JKG, KS, DHA, SD, MP, AB, MAB, RBD, FC, YZ, SQ, JH, JW and PEF performed and analysed the molecular biological, biochemical and cell biological experiments. KS and PEF performed and analyzed assembly/disassembly assay. KS and PEF performed and analyzed electron microscopy experiments, SEC and dot-blot assay. PF and DK performed and analysed the permeabilization assay. LMN, AV and SFL performed and analysed the single molecule imaging experiments. KS, ME, and PEF performed and analysed the biophysical binding assay. PSGH and GCB conceived and directed the experiments and acquired funding. All authors contributed to the writing of the manuscript.

---

**Supplementary Table 1.**
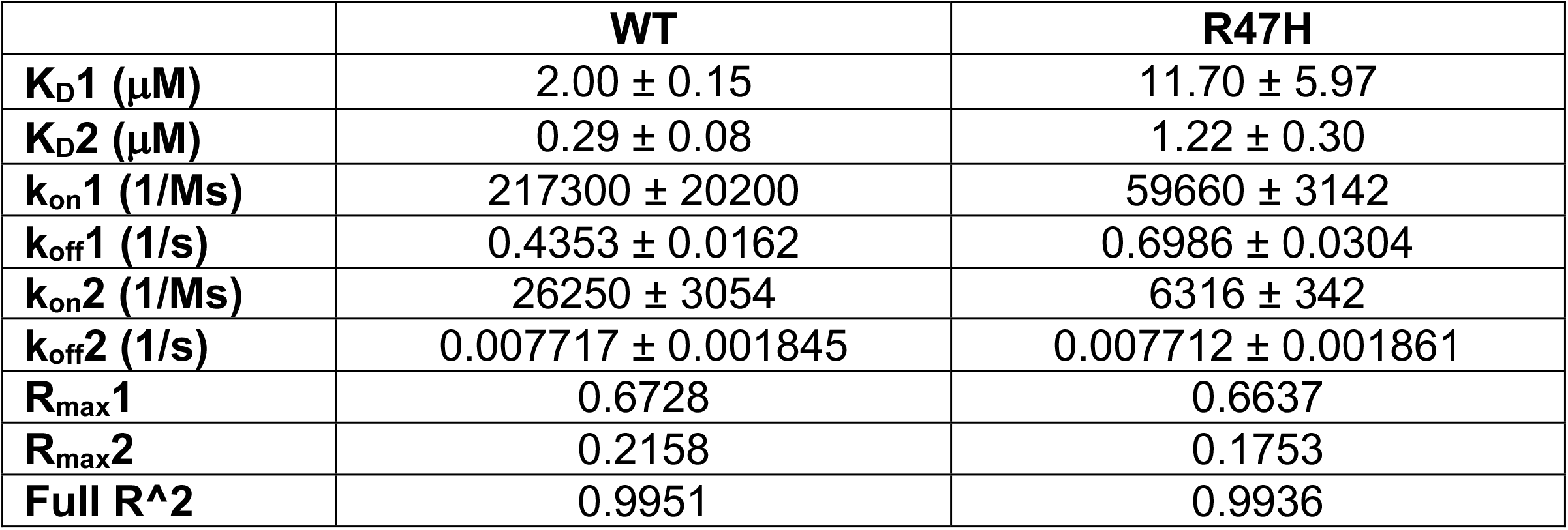
Dissociation constants, on and off rates for WT and R47H sTREM2 binding to Aβ oligomers. Fitted parameters for Supplementary Figures 6iii and iv.

**Supplementary Figure 1.**
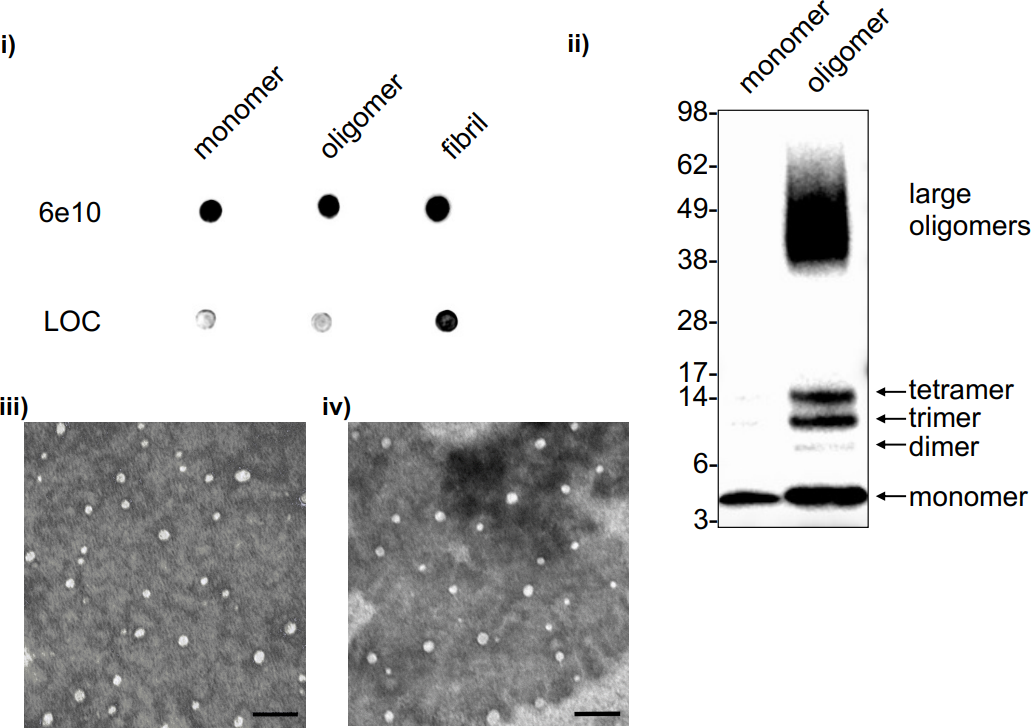
Characterization of Aβ species, i & ii): Characterization of Aβ species. Aβ42 monomers, oligomers and fibrils were prepared as previously described. i) 20 ng aliquots of Aβ monomers, oligomers and fibrils were applied to a nitrocellulose membrane and probed with mouse anti-Aβ antibody (6E10) or with rabbit anti-Amyloid fibrils antibody (LOC). ii) Aβ monomers and oligomers were identified by Western blot analysis. Aβ were separated by SDS-PAGE on a 4-12% NuPAGE bis-Tris gel and probed with anti-Aβ antibody (6E10). iii & iv) Characterization of Aβ species used in BLI experiments. Transmission electron microscopy (TEM) was performed using undiluted biotinylated Aβ (100%) or a mixture of biotinylated Aβ (10%) and unlabelled Aβ (90%) which were diluted to final concentrations of 1.1 μM. These TEM samples (10 μl) were applied to carbonate coated grids and negatively stained with 1% phosphotungstic acid (PTA). TEM micrographs indicating the globular morphology of the Aβ oligomers were obtained on a Hitachi H-7000 operated at 75kV. Scale bars are 100 nm.

**Supplementary Figure 2.**
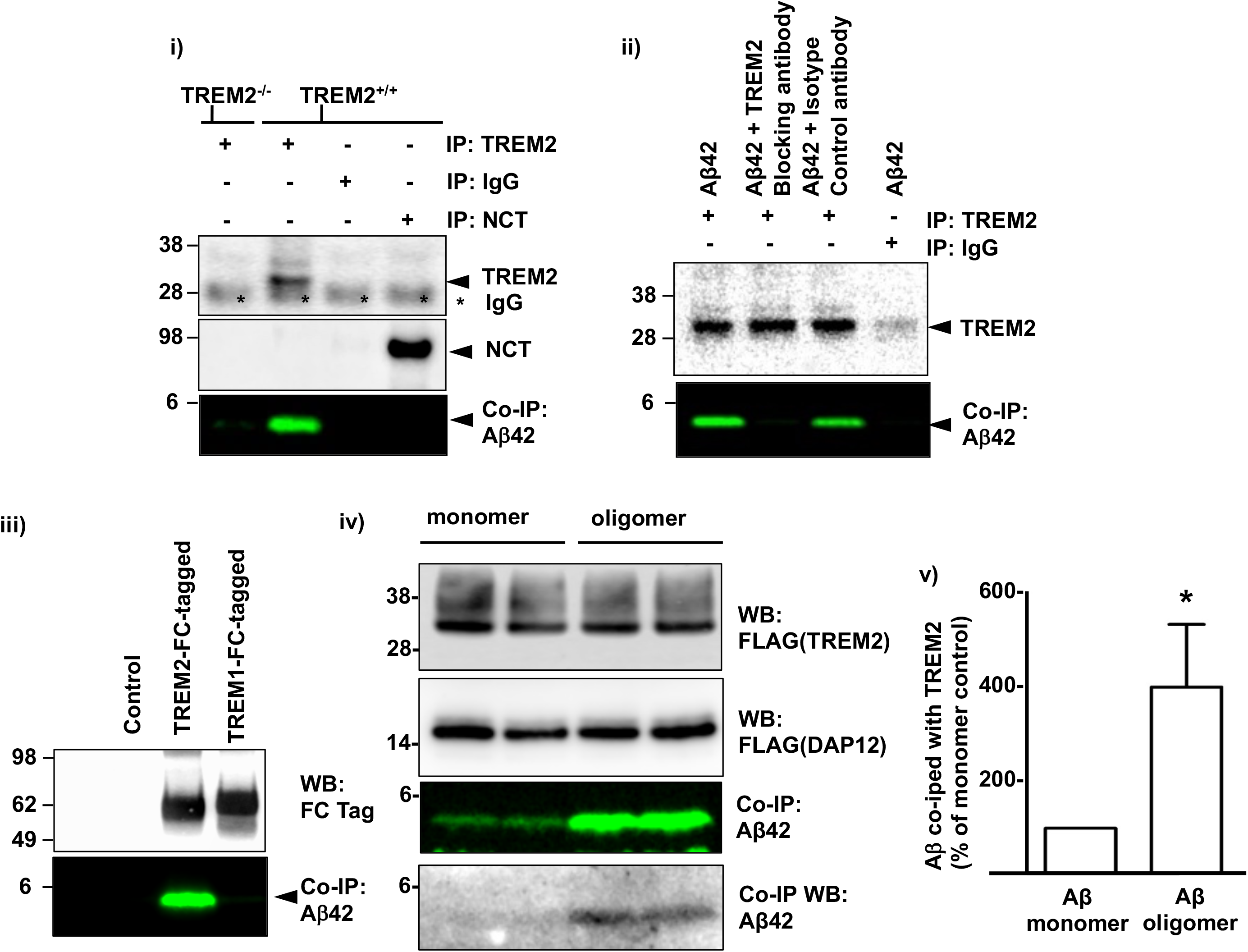
Full-length TREM2 binds Aβ oligomers but not monomers. **i)** Primary microglia were incubated with 100 nM HiLyte Fluor 647-labelled Aβ42 at 4°C for 1 hour, then lysed, immunoprecipitated with anti-TREM2 antibodies, and examined by SDS-PAGE and western blotting. Aβ oligomers co-immunoprecipitate with endogenous TREM2 in primary microglia (lane 2), but not with other natively endogenous Type I glycoproteins, such as nicastrin (lane 4) or with isotype control antibody (lane 3). TREM2 knockout microglia failed to co-immunoprecipitate Aβ (lane 1). **ii)** This interaction was blocked by preincubation with monoclonal anti-TREM2 blocking antibody (lane 2), but not by isotype control antibodies (lane 3). **iii)**: Aβ co-precipitated with recombinant FC-TREM2 (lane 2) but not FC-TREM1 (lane 3). **iv)** Aβ oligomers bind to TREM2 significantly better than monomers. HeLa cells transfected with DAP12-FLAG plus TREM2-FLAG were co-incubated with 100 nM freshly prepared monomeric or oligomeric HiLyte Fluor 647-labelled Aβ at 4°C for 1 hour. Prior to incubation, the concentration of monomer/oligomer were adjusted and tested according to fluorescence dot exposure to ensure total Aβ concentrations were the same in the working solutions of monomeric and oligomeric Aβ preparations. Cell membrane lysates were then immunoprecipitated with anti-hTREM2 antibody. The IP products were western blotted and probed with the indicated antibody. **v)** Aβ oligomers bind to TREM2 significantly better than monomers. Quantification of iv) with n = 4 replicates in 2 independent experiments; p = 0.03 Mann-Whitney U test). Error bars = SEM.

**Supplementary Figure 3.**
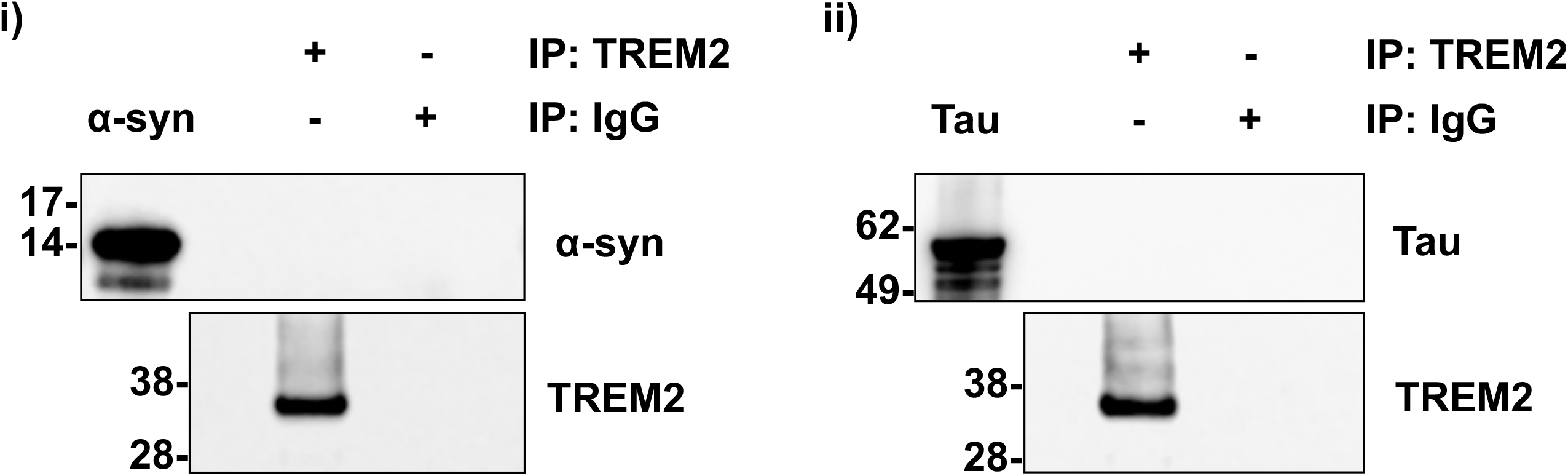
TREM2 does not bind α-synuclein or Tau oligomers. Primary microglia were incubated with 100 nM oligomeric α-synuclein or Tau protein at 4°C for 1 hour. Cell membrane lysates were then immunoprecipitated with human TREM2 antibody or control IgG. The IP products were probed with the indicated antibody.

**Supplementary Fig. 4.**
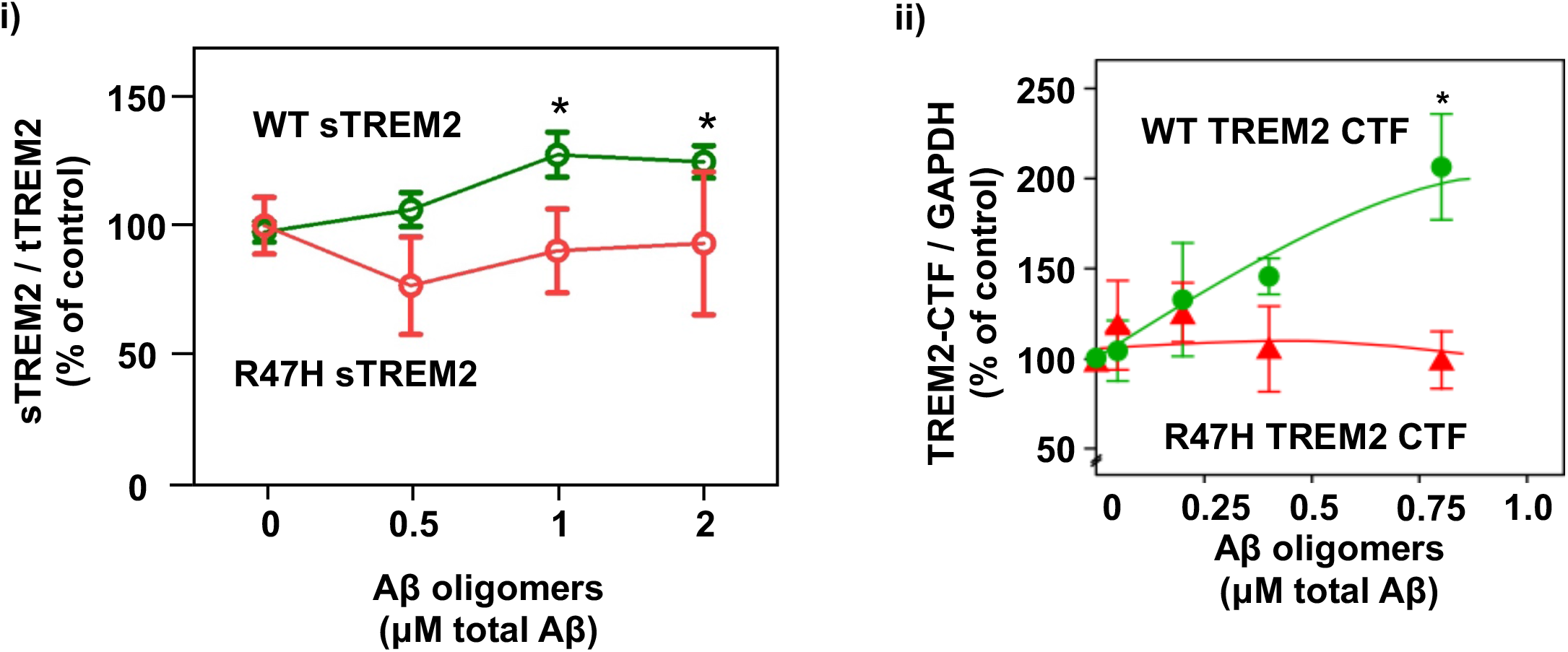
Aβ oligomers induce sTREM2 release and TREM2-CTF accumulation in cells expressing wild-type (but not R47H) TREM2. **i)** In primary murine microglia, endogenously expressed wild type TREM2 (WT-TREM2, green line) showed dose-dependent increase in shedding of soluble sTREM2 extracellular domain (green line). Aβ oligomers do not induce significant TREM2 endoproteolysis in microglia from homozygous CRISPR-Cas9 engineered R47H TREM2. Error bars = SEM; *= p <0.05, n=4 independent experiments with ≥10 replications each; one-way ANOVA with Tukey’s multiple comparisons post-test. **ii)** TREM2 membrane-bound C-terminal fragment was assayed (by western blots, Supplementary Fig. 5) 16 hours after addition of Aβ oligomers to HEK293 cells overexpressing wild-type or R47H TREM2.

**Supplementary Figure 5.**
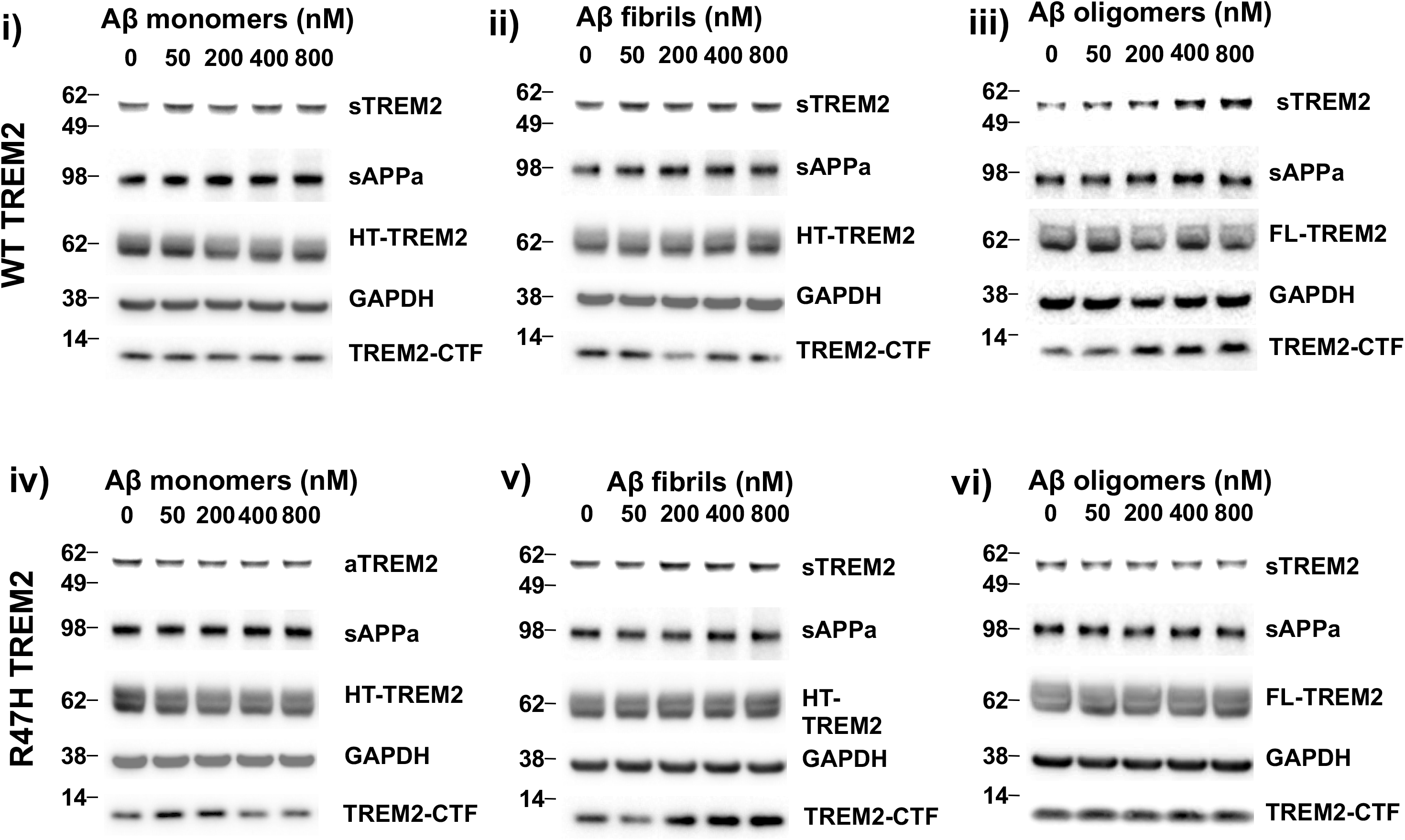
Aβ monomers and fibrils fail to trigger TREM2 cleavage, but Aβ oligomers trigger TREM2 cleavage in wild-type but not R47H TREM2-expressing cells. Aβ monomers (i & iv) fibrils (ii & v) and oligomers (iii & vi) were added at the indicated concentrations to HEK293 cells co-expressing DAP12 and either wild-type (WT, I, ii & iii) or R47H TREM2 (iv, v & vi), both HaloTagged (HT-TREM2). TREM2 membrane-bound C-terminal fragment was assayed in cell lysates and sTREM2 from the culture medium 16 hours after Aβ addition. Representative western blots shown.

**Supplementary Figure 6.**
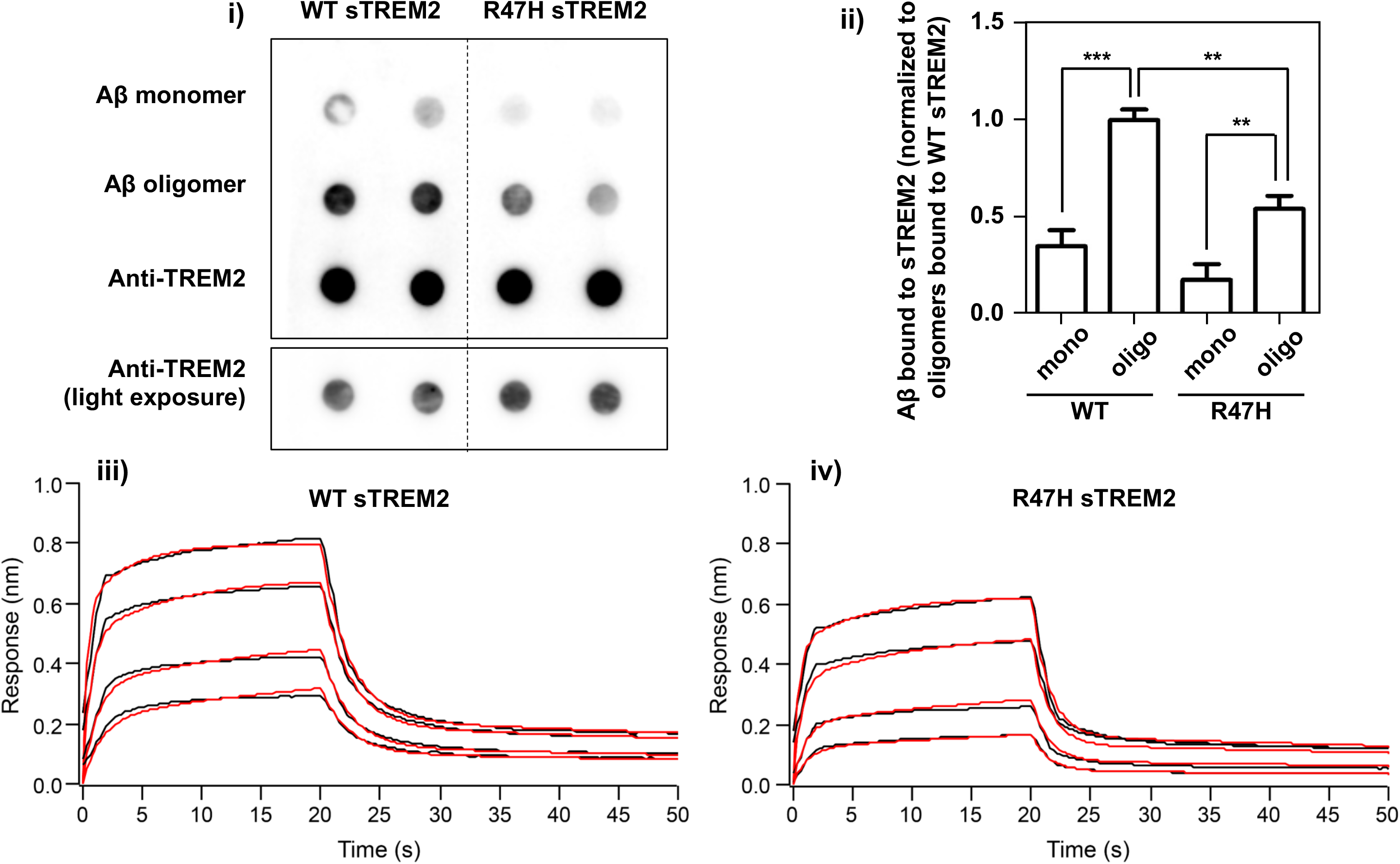
Wild-type (WT) sTREM2 binds Aβ oligomers more than Aβ monomers, and R47H sTREM2 binds both less. **i)** Dot Blot membrane strips spotted with Aβ monomers, oligomers and anti-TREM2 antibody were probed with WT sTREM2-His (left) or R47H sTREM2-His (right). Lower left panel is a lighter exposure of the anti-TREM2 antibody showing that similar amounts of sTREM2 were used. **ii)** Quantification of sTREM2 bound to Aβ. Error bars = SEM; **=p<0.01, ***=p<0.001, n=3 independent experiments with 6 replications; one-way ANOVA with Tukey’s post-hoc multiple comparisons test. **iii)** and **iv)** Bio-Layer Interferometry studies reveal a 2-state model for Aβ-binding to sTREM2. Curves of iii) WT sTREM2 or iv) R47H sTREM2 binding to Aβ oligomers at four different concentrations,. Black lines indicate experimental data. Red lines indicate fitted curves from the 2:1 model. The fitted parameters are shown in Table 1.

**Supplementary Figure 7.**
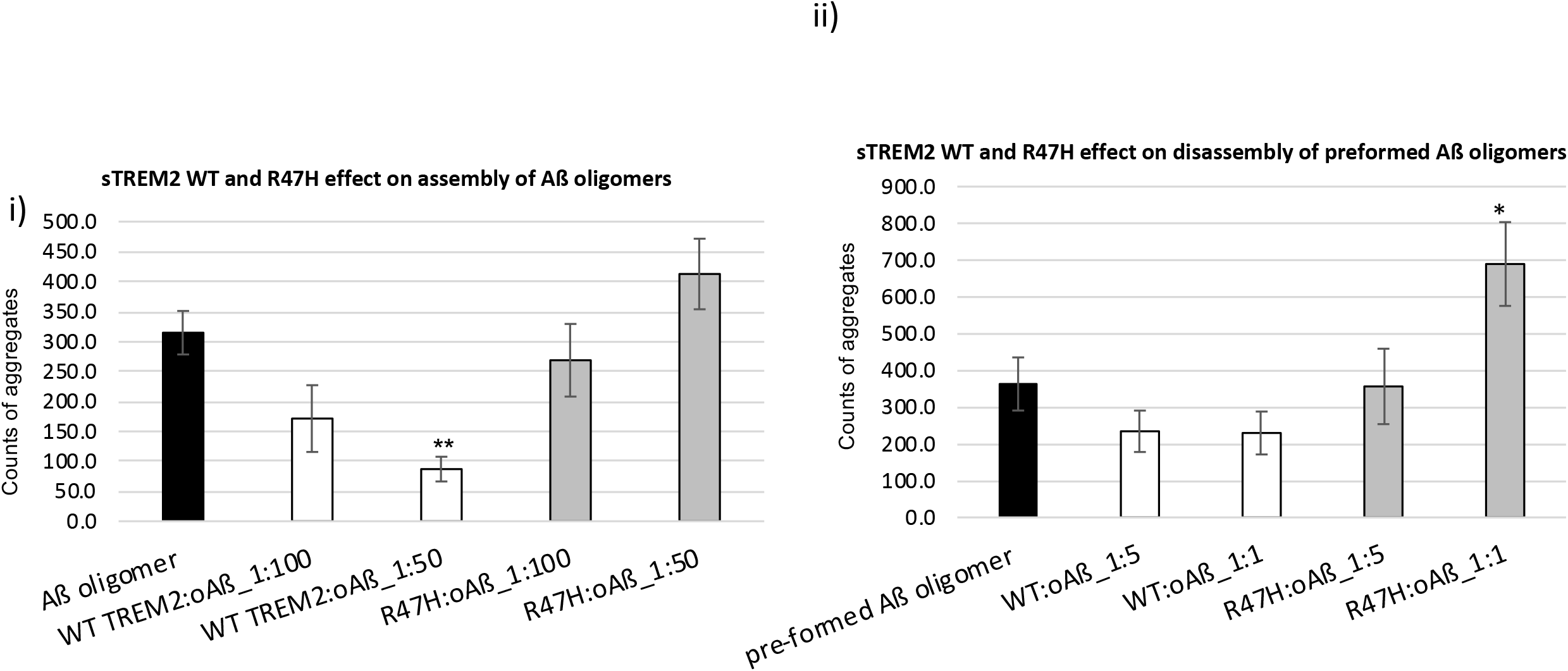
WT sTREM2 decreases Aβ oligomer assembly, whereas R47H sTREM2 increases Aβ oligomer disassembly. **i)** Aβ was oligomerized in the presence of absence of WT or R47H sTREM2 (sTREM2 1:50 or 1:100 molar ratio to Aβ). After by transmission electron microscopy imaging (representative images in Fig 2), the numbers of Aβ aggregates (oligomers) were counted. **ii)** Aβ was oligomerized, and then incubated in the presence of absence of WT or R47H sTREM2 (sTREM2 1:5 or 1:1 molar ratio to Aβ). After imaging (representative images in Fig 2), the numbers of Aβ aggregates (oligomers) were counted. Error bars represent SD. Statistical analysis was performed using one-way ANOVA followed by Bonferroni’s multiple comparison test (n=3, \**p*<0.05, \*\**p*<0.01 vs Aβ oligomer alone).

**Supplementary Figure 8.**
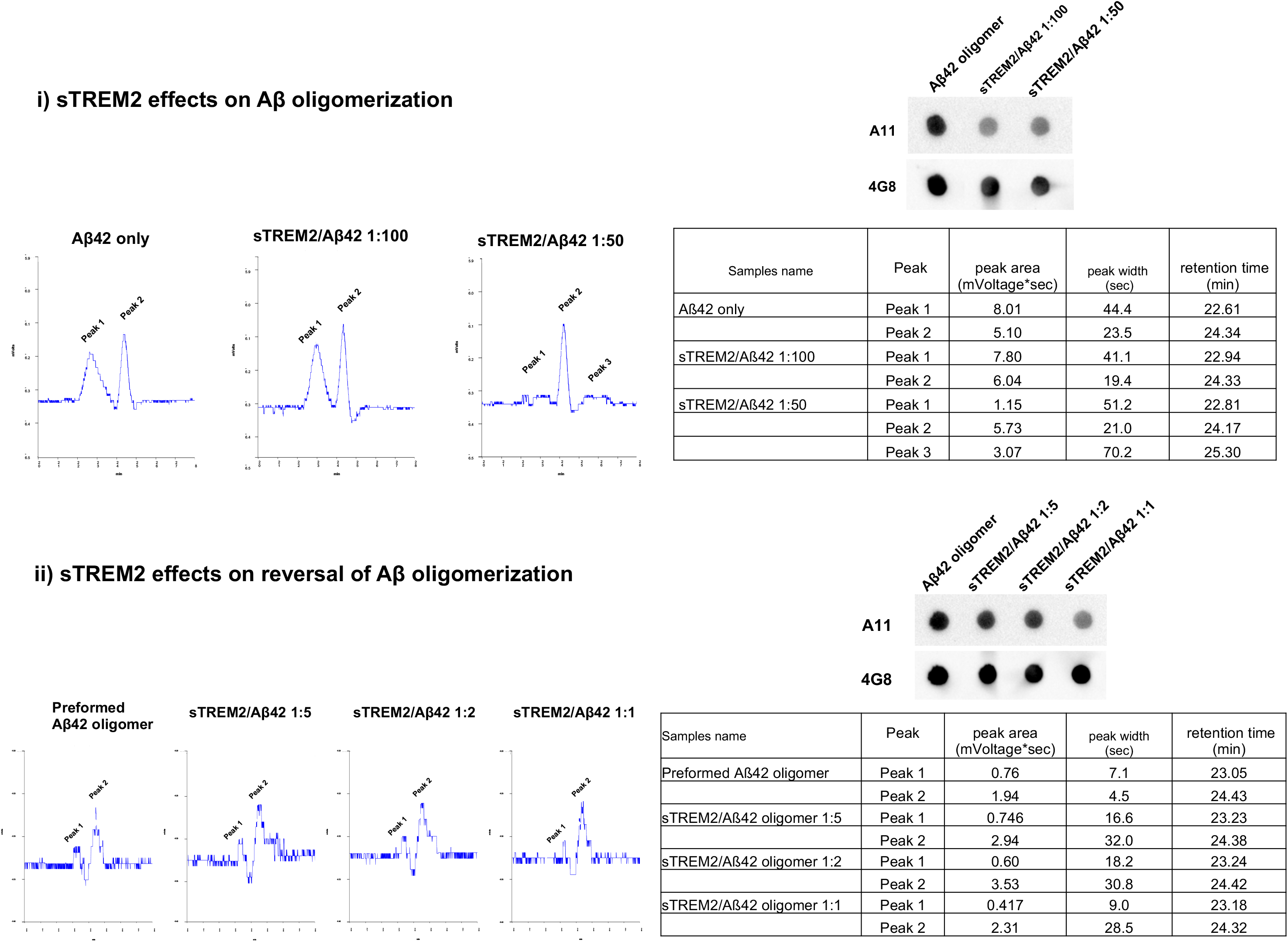
Wild-type sTREM2 inhibits oligomerisation of monomeric Aβ42, and disaggregates preformed Aβ oligomers. The same Aβ preparations examined in Figure 2i by transmission electron microscopy were also investigated by HPLC-SEC (left panels and table). In both i) Aβ oligomer formation and ii) Aβ oligomer disaggregation experiments, sTREM2 reduced the area under “Peak 1”, and increased the area under “Peak 2” and in some instances caused the appearance of a “Peak 3”. The same Aβ preparations were also examined by dot blot hybridisation against the A11 anti-oligomer antibody and the 4G8 anti-Aβ antibody (right panels). In good agreement with both the TEM and the HPLC-SEC studies, the presence of sTREM2 was associated with reductions in A11 immunoreactive oligomeric Aβ species.

**Supplementary Figure 9.**
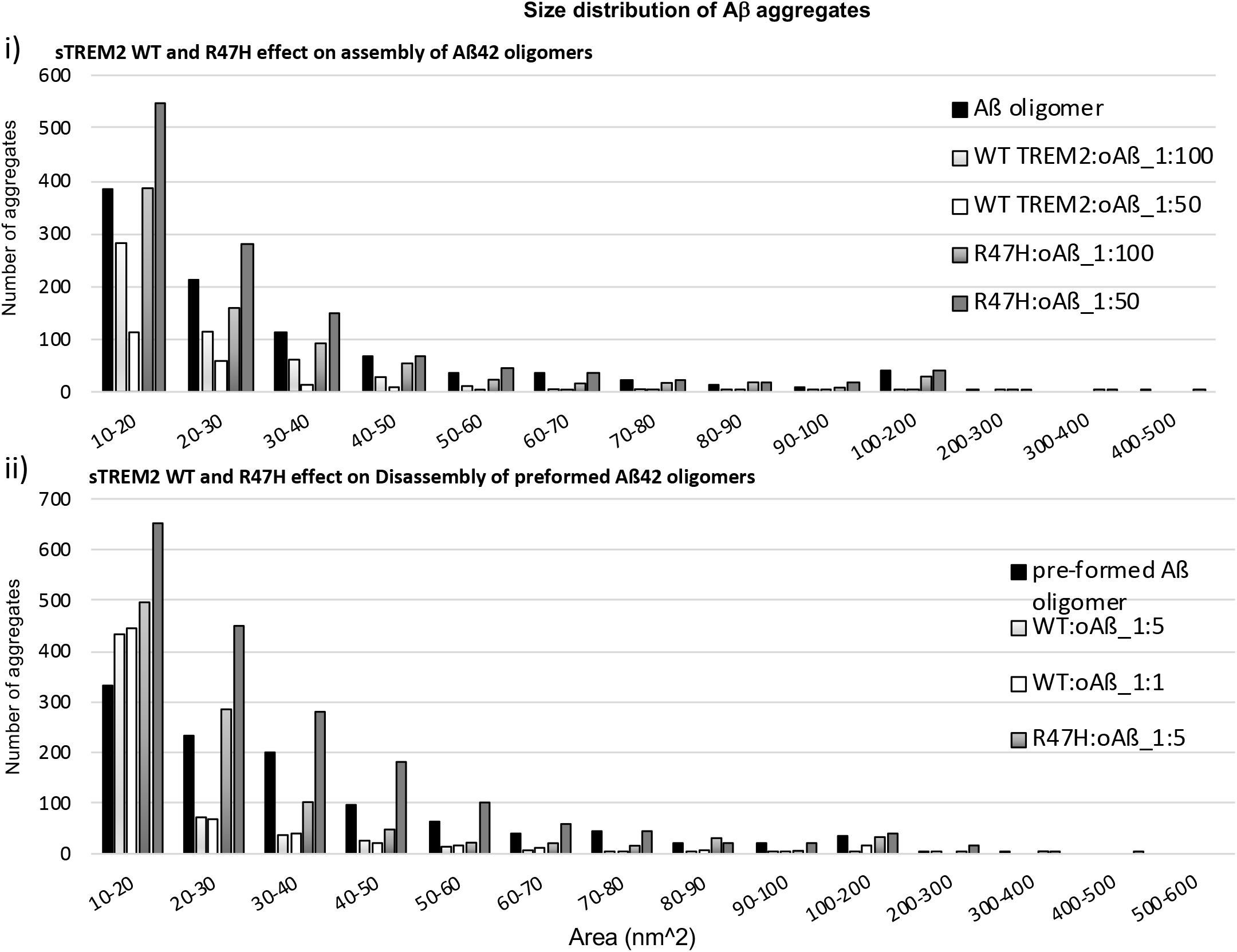
WT sTREM2 preferentially inhibits the production of large Aβ oligomers, and disaggregates large Aβ oligomers into small oligomers; R47H sTREM2 increases the number of small Aβ oligomers. **i)** Quantification of the size distribution of Aβ aggregates (oligomers) produced by oligomerizing Aβ in the presence or absence of WT or R47H sTREM2 at the indicated molar ratios. **ii)** Quantification of the size distribution of Aβ aggregates (oligomers) resulting from incubation of preformed Aβ oligomers ± WT or R47H sTREM2 at the indicated molar ratios. Representative TEM images in Fig 2. Data from three independent TEM images.

**Supplementary Figure 10.**
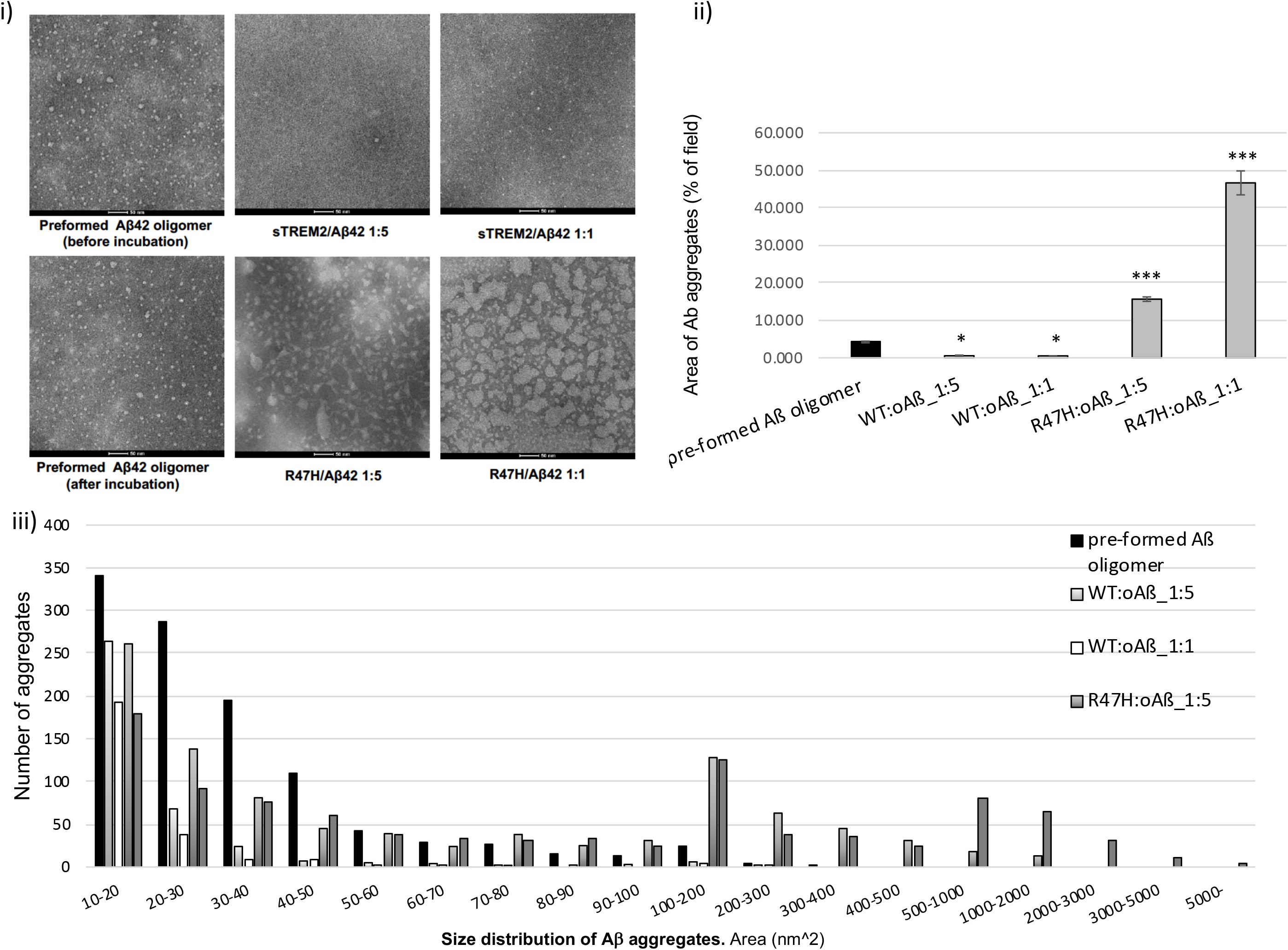
At low concentrations of Aβ, wild-type sTREM2 dissolves Aβ oligomers, whereas R47H sTREM2 induces very large Aβ aggregates: total area and size distribution of Aβ aggregates. Preformed Aβ oligomers were incubated at 100 nM Aβ (monomer equivalent) ± 20 or 100 nM wild-type or R47H sTREM2 for 30 mins at 37°C, then TEM imaged. i) Representative TEM images. ii) Quantification of total area of Aβ aggregates. Error bars represent SD. Statistical analysis was performed using one-way ANOVA followed by Bonferroni’s multiple comparison test (n=4, \**p*<0.05, \*\*\**p*<0.001 Aβ oligomer alone). iii) Size distribution of the aggregates on from three independent TEM images.

**Supplementary Figure 11.**
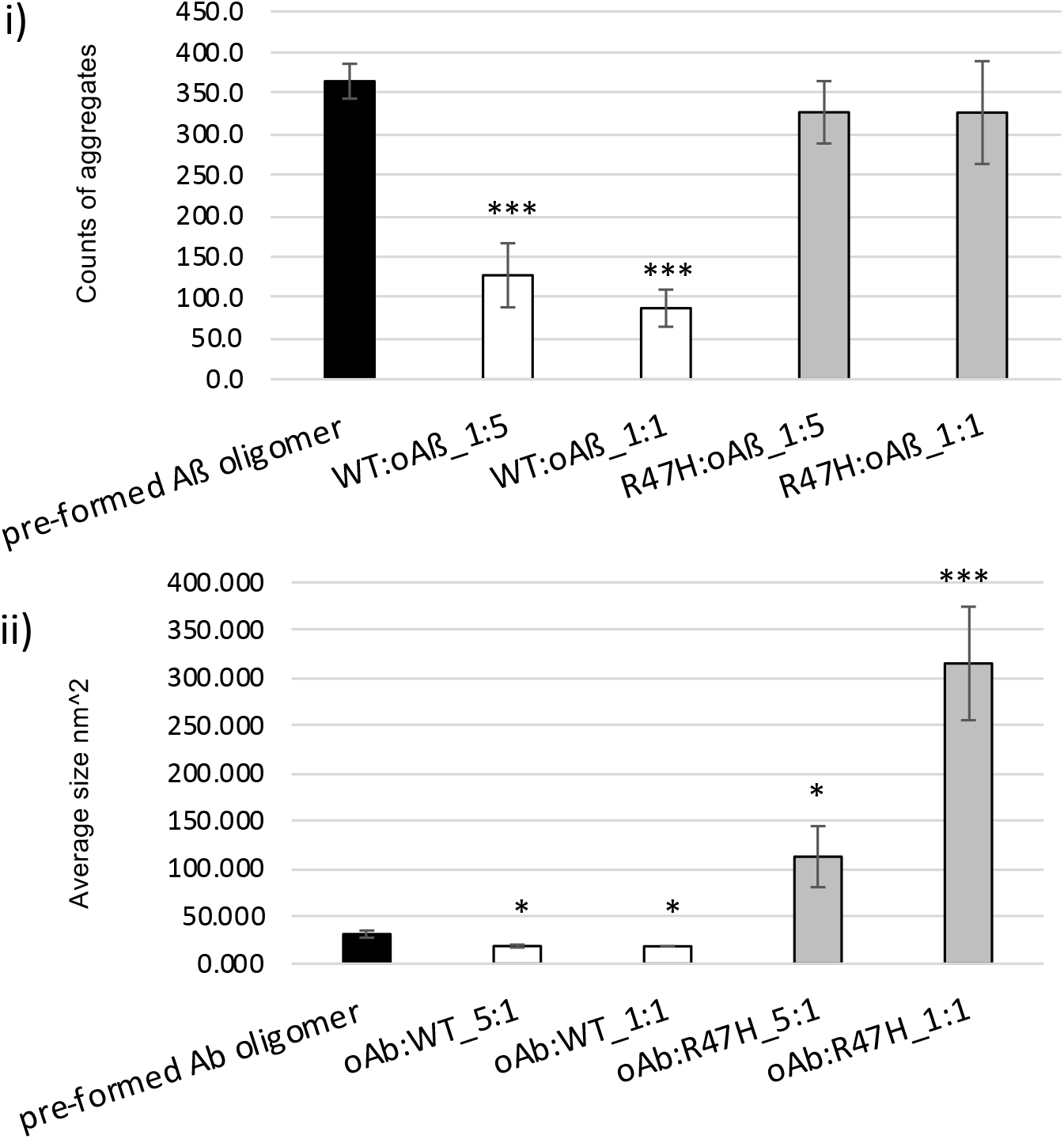
At low concentrations of Aβ, wild-type sTREM2 dissolves Aβ oligomers, whereas R47H sTREM2 induces very large Aβ aggregates: numbers and sizes of Aβ aggregates. Further analysis of the aggregates of Supplementary Fig. 10. Preformed Aβ oligomers were incubated at 100 nM Aβ (monomer equivalent) ± 20 or 100 nM wild-type or R47H sTREM2 for 30 mins, then TEM imaged. i) Number of Aβ aggregates. Ii) Average size (area) of each Aβ aggregate. Error bars represent SD. Statistical analysis was performed using one-way ANOVA followed by Bonferroni’s multiple comparison test (n=4, \**p*<0.05, \*\*\**p*<0.001 Aβ oligomer alone).

**Supplementary Figure 12.**
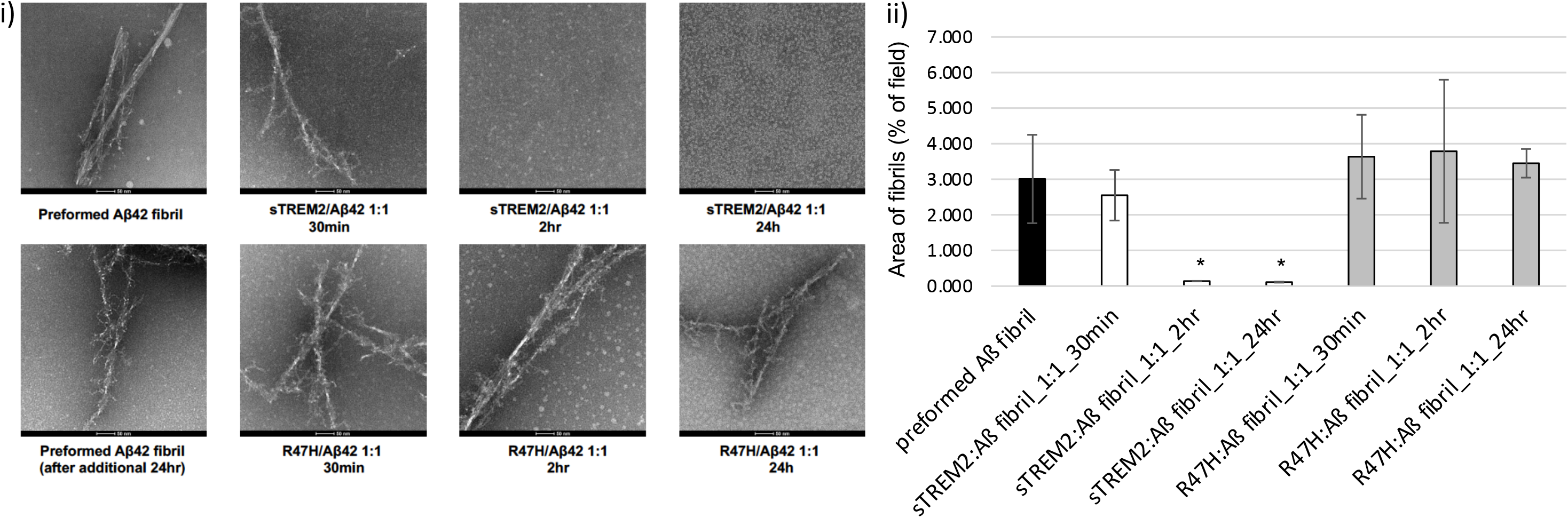
WT sTREM2 disaggregates preformed Aβ fibrils. Time course for experiment depicted in Fig. 3. **i)** Preformed Aβ fibrils were treated ± WT or R47H sTREM2 (at a 1:1 molar ratio) for 30min, 2hrs or 24 hrs. Negative-stain TEM revealed that WT sTREM2 dissociated Aβ fibrils after 2 hour incubation but R47H did not. **ii)** Quantification of the area of Aβ fibrils. Error bars represent SD. Statistical analysis was performed using one-way ANOVA followed by Bonferroni’s multiple comparison test (n=3-8,\**p*<0.05 vs pre-formed Aβ fibril).

**Supplementary Figure 13.**
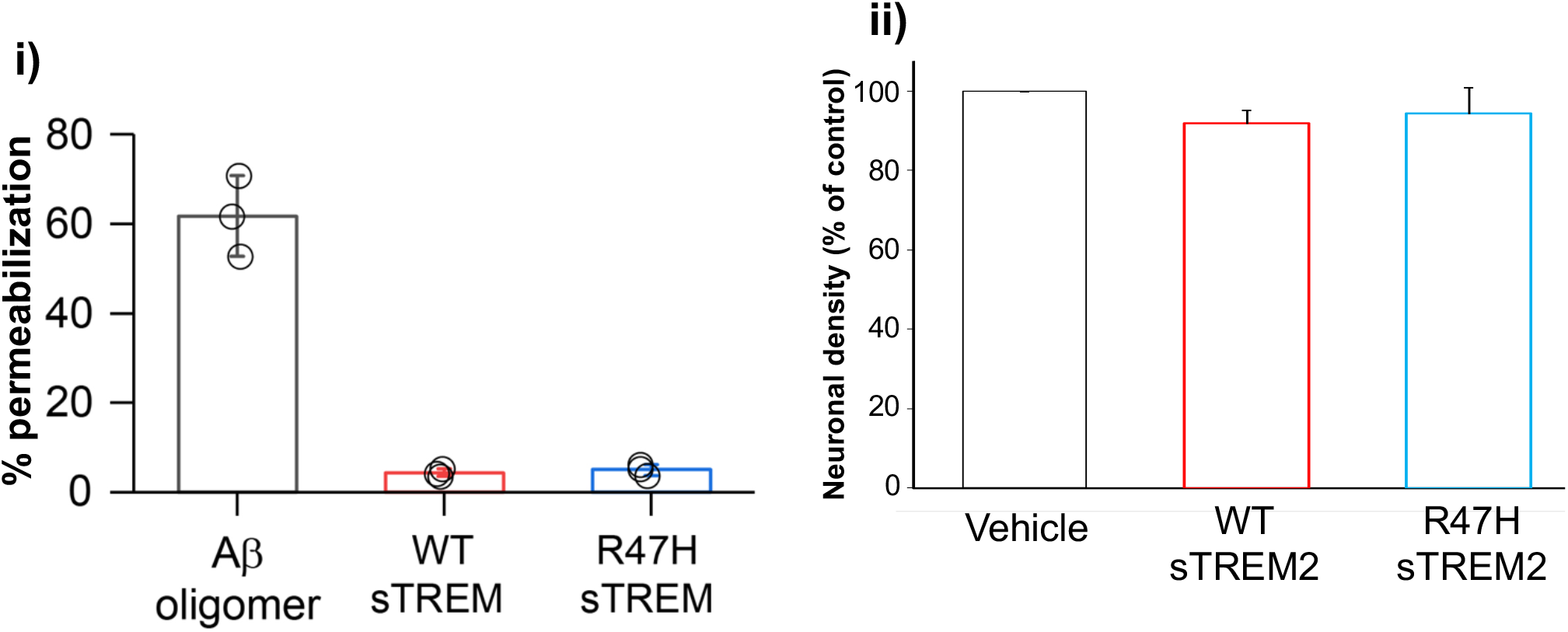
In the absence of Aβ wild-type and R47H sTREM2 have no effect on i) membrane permeability or ii) neuronal density. **i**) 1 μM Aβ, WT sTREM2 or R47H sTREM2 were incubated (separately) for 6 hours and diluted to 200 nM before the membrane-permeabilization assay was performed. Error bars = SEM; n=3 independent experiments. ii) Mixed neuronal-glial co-cultures were treated with either: vehicle, 40 nM wild-type sTREM2 or 40 nM R47H sTREM2, and 3 days later neuronal density was counted. Error bars = SEM; n=4 independent experiments on different cell cultures.

## References

1. Wong CW, Quaranta V, Glenner GG. Neuritic plaques and cerebrovascular amyloid in Alzheimer disease are antigenically related. Proc Natl Acad Sci U S A. 1985;82:8729–32.

2. Edwards FA. A Unifying Hypothesis for Alzheimer’s Disease: From Plaques to Neurodegeneration. Trends Neurosci. 2019;42:310–322. doi: 10.1016/j.tins.2019.03.003

3. Yuan P, Condello C, Keene CD, Wang Y, Bird TD, Paul SM, Luo W, Colonna M, Baddeley D, Grutzendler J. TREM2 Haplodeficiency in Mice and Humans Impairs the Microglia Barrier Function Leading to Decreased Amyloid Compaction and Severe Axonal Dystrophy. Neuron. 2016;90:724–39. doi: 10.1016/j.neuron.2016.05.003

4. Guerreiro R et al. TREM2 variants in Alzheimer’s disease. N Engl J Med 2013;368, 117–127, doi:10.1056/NEJMoa1211851.

5. Jonsson T et al. Variant of TREM2 associated with the risk of Alzheimer’s disease. N Engl J Med. 2013;368, 107–116, doi:10.1056/NEJMoa1211103.

6. Sims R et al. Rare coding variants in PLCG2, ABI3, and TREM2 implicate microglial-mediated innate immunity in Alzheimer’s disease. Nat Genet 2017;49, 1373–1384, doi:10.1038/ng.3916.

7. Wunderlich P, Glebov K, Kemmerling N, Tien NT, Neumann H, Walter J. Sequential proteolytic processing of the triggering receptor expressed on myeloid cells-2 (TREM2) protein by ectodomain shedding and γ-secretase-dependent intramembranous cleavage. J Biol Chem. 2013;288:33027–36. doi: 10.1074/jbc.M113.517540

8. Thornton P, Sevalle J, Deery MJ, Fraser G, Zhou Y, Ståhl S, Franssen EH, Dodd RB, Qamar S, Gomez Perez-Nievas B, Nicol LS, Eketjäll S, Revell J, Jones C, Billinton A, St George-Hyslop PH, Chessell I, Crowther DC. TREM2 shedding by cleavage at the H157-S158 bond is accelerated for the Alzheimer’s disease-associated H157Y variant. EMBO Mol Med. 2017;9:1366–1378. doi: 10.15252/emmm.201707673

9. Schlepckow K, Kleinberger G, Fukumori A, Feederle R, Lichtenthaler SF, Steiner H, Haass C. An Alzheimer-associated TREM2 variant occurs at the ADAM cleavage site and affects shedding and phagocytic function. EMBO Mol Med. 2017;9:1356–1365. doi: 10.15252/emmm.201707672

10. Suárez-Calvet M et al. Early increase of CSF sTREM2 in Alzheimer’s disease is associated with tau related-neurodegeneration but not with amyloid-β pathology. Mol Neurodegener. 2019;14:1. doi: 10.1186/s13024-018-0301-5

11. Suarez-Calvet M et al. Early changes in CSF sTREM2 in dominantly inherited Alzheimer’s disease occur after amyloid deposition and neuronal injury. Sci Transl Med 2016;8:369ra178, doi:10.1126/scitranslmed.aag1767.

12. Suarez-Calvet M et al. sTREM2 cerebrospinal fluid levels are a potential biomarker for microglia activity in early-stage Alzheimer’s disease and associate with neuronal injury markers. EMBO Mol Med 2016;8:466–476. doi:10.15252/emmm.201506123.

13. Zhao Y et al. TREM2 Is a Receptor for beta-Amyloid that Mediates Microglial Function. Neuron 2018;97:1023–1031. doi:10.1016/j.neuron.2018.01.031.

14. Zhong L, Wang Z, Wang D, Wang Z, Martens YA, Wu L, Xu Y, Wang K, Li J, Huang R, Can D, Xu H, Bu G, Chen XF. Amyloid-beta modulates microglial responses by binding to the triggering receptor expressed on myeloid cells 2 (TREM2). Mol Neurodegener. 2018;13:15.

15. Lessard CB, Malnik SL, Zhou Y, Ladd TB, Cruz PE, Ran Y, Mahan TE, Chakrabaty P, Holtzman DM, Ulrich JD, Colonna M, Golde TE. High-affinity interactions and signal transduction between Aβ oligomers and TREM2. EMBO Mol Med. 2018;10:e9027.

16. Zhong L, Xu Y, Zhuo R, Wang T, Wang K, Huang R, Wang D, Gao Y, Zhu Y, Sheng X, Chen K, Wang N, Zhu L, Can D, Marten Y, Shinohara M, Liu CC, Du D, Sun H, Wen L, Xu H, Bu G, Chen XF. Soluble TREM2 ameliorates pathological phenotypes by modulating microglial functions in an Alzheimer’s disease model. Nat Commun. 2019;10:1365.

17. Parhizkar S, Arzberger T, Brendel M, Kleinberger G, Deussing M, Focke C, Nuscher B, Xiong M, Ghasemigharagoz A, Katzmarski N, Krasemann S, Lichtenthaler SF, Müller SA, Colombo A, Monasor LS, Tahirovic S, Herms J, Willem M, Pettkus N, Butovsky O, Bartenstein P, Edbauer D, Rominger A, Ertürk A, Grathwohl SA, Neher JJ, Holtzman DM, Meyer-Luehmann M, Haass C. Loss of TREM2 function increases amyloid seeding but reduces plaque-associated ApoE. Nat Neurosci. 2019;22:191–204.

18. Wang Y, Ulland TK, Ulrich JD, Song W, Tzaferis JA, Hole JT, Yuan P, Mahan TE, Shi Y, Gilfillan S, Cella M, Grutzendler J, DeMattos RB, Cirrito JR, Holtzman DM, Colonna M. TREM2-mediated early microglial response limits diffusion and toxicity of amyloid plaques. J Exp Med. 2016;213:667–75. doi: 10.1084/jem.20151948

19. Ma LZ, Tan L, Bi YL, Shen XN, Xu W, Ma YH, Li HQ, Dong Q, Yu JT. Dynamic changes of CSF sTREM2 in preclinical Alzheimer’s disease: the CABLE study. Mol Neurodegener. 2020;15:25. doi: 10.1186/s13024-020-00374-8

20. Ewers M et al. Increased soluble TREM2 in cerebrospinal fluid is associated with reduced cognitive and clinical decline in Alzheimer’s disease. Sci Transl Med. 2019;11:507. doi: 10.1126/scitranslmed.aav6221

21. Franzmeier N, Suárez-Calvet M, Frontzkowski L, Moore A, Hohman TJ, Morenas-Rodriguez E, Nuscher B, Shaw L, Trojanowski JQ, Dichgans M, Kleinberger G, Haass C, Ewers M; Alzheimer’s Disease Neuroimaging Initiative (ADNI). Higher CSF sTREM2 attenuates ApoE4-related risk for cognitive decline and neurodegeneration. Mol Neurodegener. 2020;15:57. doi: 10.1186/s13024-020-00407-2

22. Ewers M, Biechele G, Suárez-Calvet M, Sacher C, Blume T, Morenas-Rodriguez E, Deming Y, Piccio L, Cruchaga C, Kleinberger G, Shaw L, Trojanowski JQ, Herms J, Dichgans M; Alzheimer’s Disease Neuroimaging Initiative (ADNI), Brendel M, Haass C, Franzmeier N. Higher CSF sTREM2 and microglia activation are associated with slower rates of beta-amyloid accumulation. EMBO Mol Med. 2020;12:e12308. doi: 10.15252/emmm.202012308

23. Horrocks MH et al. Single-Molecule Imaging of Individual Amyloid Protein Aggregates in Human Biofluids. ACS Chem Neurosci 2016;7:399–406. doi:10.1021/acschemneuro.5b00324.

24. Concepcion J et al. Label-free detection of biomolecular interactions using Bio-Layer Interferometry for kinetic characterization. Comb Chem High Throughput Screen 2009;12:791–800.

25. Harper JD, Wong SS, Lieber CM, Lansbury PT Jr. Assembly of A beta amyloid protofibrils: an in vitro model for a possible early event in Alzheimer’s disease. Biochemistry. 1999 Jul 13;38(28):8972–80.

26. Nichols MR1, Moss MA, Reed DK, Cratic-McDaniel S, Hoh JH, Rosenberry TL. Amyloid-beta protofibrils differ from amyloid-beta aggregates induced in dilute hexafluoroisopropanol in stability and morphology. J Biol Chem. 2005;280:2471–80.

27. De S, Whiten DR, Ruggeri FS, Hughes C, Rodrigues M, Sideris DI, Taylor CG, Aprile FA, Muyldermans S, Knowles TPJ, Vendruscolo M, Bryant C, Blennow K, Skoog I, Kern S, Zetterberg H, Klenerman D. Soluble aggregates present in cerebrospinal fluid change in size and mechanism of toxicity during Alzheimer’s disease progression. Acta Neuropathol Commun. 2019;7:120.

28. De S, Wirthensohn DC, Flagmeier P, Hughes C, Aprile FA, Ruggeri FS, Whiten DR, Emin D, Xia Z, Varela JA, Sormanni P, Kundel F, Knowles TPJ, Dobson CM, Bryant C, Vendruscolo M, Klenerman D. Different soluble aggregates of Aβ42 can give rise to cellular toxicity through different mechanisms. Nat Commun. 2019;10:1541.

29. Neniskyte U, Neher JJ, Brown GC. Neuronal death induced by nanomolar amyloid β is mediated by primary phagocytosis of neurons by microglia. J Biol Chem. 2011;286:39904–13.

30. Piccio L, Deming Y, Del-Águila JL, Ghezzi L, Holtzman DM, Fagan AM, Fenoglio C, Galimberti D, Borroni B, Cruchaga C. Cerebrospinal fluid soluble TREM2 is higher in Alzheimer disease and associated with mutation status. Acta Neuropathol. 2016;131:925–33.

31. Yang T, Xu H, Walsh DM, Selkoe DJ. Large Soluble Oligomers of Amyloid β-Protein from Alzheimer Brain Are Far Less Neuroactive Than the Smaller Oligomers to Which They Dissociate. J Neurosci 2017; 35:152–163.

32. Iljina M, Hong L, Horrocks MH, Ludtmann MH, Choi ML, Hughes CD, Ruggeri FS, Guilliams T, Buell AK, Lee JE, Gandhi S, Lee SF, Bryant CE, Vendruscolo M, Knowles TPJ, Dobson CM, sDe Genst E, Klenerman D. Nanobodies raised against monomeric ɑ-synuclein inhibit fibril formation and destabilize toxic oligomeric species. BMC Biol. 2017;15:57.

33. Kang SS et al. Behavioral and transcriptomic analysis of Trem2-null mice: not all knockout mice are created equal. Hum Mol Genet 2018;27:211–223. doi:10.1093/hmg/ddx366.

33. Bal-Price A, Brown GC. Inflammatory neurodegeneration mediated by nitric oxide from activated glia-inhibiting neuronal respiration, causing glutamate release and excitotoxicity. Journal of Neuroscience 2001;21:6480–6491.

34. Stine WB Jr, Dahlgren KN, Krafft GA, LaDu MJ. In vitro characterization of conditions for amyloid-beta peptide oligomerization and fibrillogenesis. J Biol Chem 2003;278:11612–11622. doi:10.1074/jbc.M210207200.

35. Drews A et al. Individual aggregates of amyloid beta induce temporary calcium influx through the cell membrane of neuronal cells. Sci Rep 2016;6:31910. doi:10.1038/srep31910.

36. Sevalle J et al. Aminopeptidase A contributes to the N-terminal truncation of amyloid beta-peptide. J Neurochem 2009;109:248–256. doi:10.1111/j.1471-4159.2009.05950.x.

37. Chia S et al. Monomeric and fibrillar α-synuclein exert opposite effects on the catalytic cycle that promotes the proliferation of Aβ42 aggregates. Proc. Natl. Acad. Sci. 2017;114:8005–8010.

38. Flagmeier P et al. Ultrasensitive measurement of Ca^2+^ influx into lipid vesicles induced by protein aggregates. Angew. Chem. Int. Ed. 2017;56:7750–7754.

